# Atypical peripheral actin band formation via overactivation of RhoA and Non-muscle myosin II in Mitofusin 2 deficient cells

**DOI:** 10.1101/2022.10.03.510661

**Authors:** Yueyang Wang, Lee D. Troughton, Fan Xu, Aritra Chatterjee, Han Zhao, Laura P. Cifuentes, Ryan B. Wagner, Jingjuan Chen, Shihuan Kuang, Daniel M. Suter, Chongli Yuan, Deva Chan, Fang Huang, Patrick W. Oakes, Qing Deng

**Affiliations:** Department of Biological Sciences, Purdue University, West Lafayette, IN 47907, USA; Cell and Molecular Physiology, Loyola University Chicago, Stritch School of Medicine, Maywood, IL 60153, USA; Weldon School of Biomedical Engineering, Purdue University, West Lafayette, IN47907, USA; Advanced Research Institute of Multidisciplinary Science, Beijing Institute of Technology, Beijing100081, China; Davidson School of Chemical Engineering, Purdue University, West Lafayette, IN47907, USA; School of Mechanical Engineering, Purdue University, West Lafayette, IN 47907, USA; Department of Animal Sciences, Purdue University, West Lafayette, IN, 47907, USA; Purdue Institute for Integrative Neuroscience, Purdue University, West Lafayette, IN, 47907, USA; Purdue Institute for Inflammation, Immunology & Infectious Disease, Purdue University, West Lafayette, IN 47907, USA; Purdue University Center for Cancer Research, Purdue University, West Lafayette, IN 47907, USA

## Abstract

Cell spreading and migration play central roles in many physiological and pathophysiological processes. We have previously shown that MFN2 regulates the migration of human neutrophil-like cells via suppressing Rac activation. Here, we show that in mouse embryonic fibroblasts, MFN2 suppresses RhoA activation and supports cell polarization. After the initial spreading period, the wild-type cells polarize and migrate, whereas the *Mfn2*^-/-^ cells maintain a circular shape. Increased cytosolic Ca^2+^ resulting from the loss of Mfn2 is directly responsible for this phenotype, which can be rescued by expressing an artificial tether to bring mitochondria and ER to close vicinity. Elevated cytosolic Ca^2+^ activates Ca^2+^/calmodulin-dependent protein kinase II, RhoA, and Myosin light-chain kinase, causing an over-activation of non-muscle myosin II and a formation of a prominent F-actin ring at the cell periphery and increased cell contractility. The formation of the peripheral actin band alters cell physics and is dependent on substrate rigidity. Our results provide a novel molecular basis to understand how MFN2 regulates distinct signaling pathways in different cells and tissue environments, which is instrumental in understanding and treating MFN2-related diseases.

## Introduction

Cell spreading and migration play central roles in many physiological and pathophysiological processes. The dynamic cytoskeletal reorganization during cell migration is primarily achieved through a delicate balance between protrusive and retractive forces. The cytoskeleton and regulatory proteins cooperate with spatial and temporal precision to organize cell contents to control protrusions, adhesion, contractility, and force transmission(Lauffenburger and Horwitz, 1996; Pollard and Borisy, 2003; Seetharaman and Etienne-Manneville, 2020). The cytoskeletal networks are controlled by master regulators, such as the small Rho GTPases (Nobes and Hall, 1995; Takai et al., 1995; Kaibuchi et al., 1999). The initial cell spreading is driven by actin polymerization promoted by Rac1 and Cdc42 to form a sheet-like protrusion that generates a pushing force at the cell’s leading edge. Subsequently, Ras homolog gene family member A (RhoA) and calcium/calmodulin (CaM)-dependent pathways modulate myosin-dependent contractile force by regulating focal adhesions (Ridley and Hall, 1992; Nobes and Hall, 1999, 1995) and inducing the formation of actin-myosin filaments, which form stress fibers (Ridley and Hall, 1992; Amano et al., 1996). Reduced activity of RhoA is necessary for spreading and migration, which facilitates cell edge extension by reducing myosin-dependent contractile forces (Wakatsuki et al., 2003). Myosin, specifically Non-muscle Myosin II (NMII), functions as a master regulator of cell stiffness, further influencing cell migration(Tee et al., 2011). Notably, in addition to generating mechanical force within a cell, NMII plays an essential role in sensing and responding to external forces applied to the cell(Vicente-Manzanares et al., 2009; Aguilar-Cuenca et al., 2014; Lamb et al., 2021).

Fibroblasts are mesenchyme-derived cells that play an essential role in supporting tissue development and repair by remodeling the extracellular matrix. Additionally, they secrete multiple growth factors and respond to migratory cues such as PDGF (Wynn, 2008). As a widely used cell model, the *in vitro* motility of fibroblasts has been extensively studied. Local Ca^2+^ pulses play critical roles in migrating cells, including fibroblasts, and the Ca^2+^ homeostasis controls the organization of the cytoskeleton spatially and temporally (Bennett and Weeds, 1986; Tsai et al., 2015; Tsai and Meyer, 2012). The intracellular Ca^2+^ signals are predominantly generated from the intracellular Ca^2+^ storage, the endoplasmic reticulum (ER), through inositol triphosphate (IP3) receptors (Clapham, 2007a; Parys and de Smedt, 2012). Calmodulin (CaM) is an essential effector protein in cells to amplify the Ca^2+^ signaling (Clapham, 2007b). The Ca^2+^/calmodulin (CaM)-dependent pathways promote the phosphorylation of the myosin light chain (MLC), promoting the formation of adhesive contacts and stress fibers(Kamm and Stull, 1985; Stull et al., 1998). In addition, Ca^2+^/CaM activates Ca^2+^/CaM kinases (CaMKs), including CaMKI, CaMKK, and CaMKII (Saneyoshi and Hayashi, 2012; Soderling, 1999; Hudmon and Schulman, 2002), each regulates actin cytoskeleton in distinct pathways(Saneyoshi and Hayashi, 2012). Notably, CaMKII bundles F-actin to remodel the cytoskeleton (Lin and Redmond, 2008; Okamoto et al., 2007; O’Leary et al., 2006) and regulates Rho GTPases, including Rac and RhoA, by phosphorylating their GEFs and GAPs (Fleming et al., 1999; Okabe et al., 2003; Tolias et al., 2005; Xie et al., 2007; Penzes et al., 2008).

Mitochondria are central cellular power stations. In addition, they regulate many physiological processes, such as maintaining intracellular Ca^2+^ homeostasis and cell migration (Denisenko et al., 2019; Campello et al., 2006; Zhao et al., 2012; Báthori et al., 2006). The Mitofusins (MFN 1 and MFN2) localize to the outer mitochondrial membrane (OMM) and form homo- or heterodimers to promote mitochondrial outer membrane tethers (Santel and Fuller, 2001; Chen et al., 2003a). Human MFN1 and MFN2 share ~80% similarity in protein sequence (Dc et al., 2004). They both contain a large, cytosolic, N-terminal GTPase domain, two coiled-coil heptad-repeat (HR) domains, and two transmembrane domains (TM) crossing the OMM. MFN1 and MFN2 have primarily overlapping functions. Overexpression of either protein in MFN1 or MFN2 null cells promotes mitochondrial fusion (Chen et al., 2003a). Knocking out either MFN1 or MFN2 leads to fragmented mitochondria in fibroblasts (Chen et al., 2003b; Cipolat et al., 2004). Structural and biochemical studies revealed the difference between MFN1 and MFN2 in catalytic GTPase activity (Ishihara et al., 2004; Li et al., 2019) and in their ability to mediate trans-organelle calcium signaling (Dorn, 2020; Naon et al., 2016a; de Brito and Scorrano, 2008c). MFN2, but not MFN1, localizes to the mitochondria-associated ER membranes (MAM)(de Brito and Scorrano, 2008c; Filadi et al., 2015a). Mfn2 ablation in various cell types results in increased distance between the ER and mitochondria and severely reduced Ca^2+^ transfer from the ER to mitochondria (de Brito and Scorrano, 2008a; Filadi et al., 2015b; Naon et al., 2016b). Investigation of MFN2’s role in human diseases has primarily focused on MFN-mediated mitochondrial fusion, trafficking, metabolism, mitophagy, and mitochondrial quality control. How MFNs regulate the cytoskeleton, however, remains unclear.

In our previous research to understand the importance of mitochondrial shape in neutrophil migration, we generated transgenic zebrafish lines with CRISPR-based neutrophil-specific knockout of mitochondrial fusion-related genes (Maianski et al., 2002; Zhou et al., 2018). Surprisingly, we noticed a phenotype specific to Mfn2 deletion: most neutrophils exited the hematopoietic tissue and circulated in the bloodstream in homeostasis. We further demonstrated that MFN2 regulates neutrophil adhesive migration and Rac activation using the human neutrophil-like differentiated HL-60 cells (Zhou et al., 2021). Although we identified an essential role for MFN2 in neutrophil adhesion and migration, it is unclear how MFN2 regulates actin cytoskeleton organization and other cellular behaviors, such as cell spreading.

Here, we used mice embryonic fibroblasts (MEFs) as a model to further characterize how MFN2 regulates cytoskeletal organization. We demonstrate that MFN2 regulates cytoskeletal organization by suppressing Rho and NMII activity. *Mfn2* depletion upregulates cytosolic calcium in MEFs, leading to RhoA and NMII overactivation and forming a prominent “Peripheral Actin Band (PAB)” structure. This PAB hampered cell adhesive migration and caused significant changes in mechanical properties, including cell stiffness and membrane tension. Together, our results provided an in-depth molecular understanding of the role of MFN2 in cytoskeleton dynamics, cell spreading, and adhesive migration, which may lead to a better understanding and treatment of MFN2-associated diseases.

## Results

### MFN2 deficiency changes cell morphology and impairs adhesive 2D random migration in MEFs

As a first step in investigating the role of MFN2 in MEF cells, we confirmed the respective protein loss in indicated cell lines by immunoblotting (Fig.1 A). We first analyzed the morphology and size of the cells in culture and found that the average cell size was reduced significantly in *Mfn2*-null MEFs (1113 μm^2^) as compared to *wt* (1854 μm^2^) and *Mfn1*-null MEFs (2385 μm^2^) (Fig. 1B). *Mfn2*-null MEFs also displayed significantly increased cell circularity (Fig. 1C). To evaluate the function of MFN2 protein in the cytoskeleton and cell migration, we seeded the cells on chamber slides and imaged them overnight using phase-contrast, time-lapse microscopy. In *Mfn2*-null MEFs, cell motility (0.23 ± 0.08 μm/min) was significantly reduced as compared to *wt* (0.53 ± 0.16 μm/min), or *Mfn1*-null MEFs (0.49 ± 0.12 μm/min) (Fig. 1F-H, Movie S1). No significant change in directionality was observed in *Mfn2*-null MEFs (Fig. 1I). During cell spreading, *wt* and *Mfn1* -null MEFs generated rapid protrusive filopodia and lamellipodia, eventually elongating to form traditional fibroblast-like shapes and began to migrate. However, the elimination of MFN2 caused significant defects in elongation, and the cells remained rounded (Fig. 1D-E). The morphological differences became obvious during the spreading process, especially after 20 mins. *Mfn2*-null MEFs only extended round membrane ruffles but did not simultaneously form multiple short lamellae separated by concave edges. Immunofluorescence also revealed striking differences in actin stress fiber organization. Both *wt* and *Mfn1*-null MEFs displayed parallel stress fibers in the cell body, while *Mfn2*-null MEFs contained an enrichment in actin filaments in the cell cortex with reduced stress fibers at the center of the cells (Fig. S1A). To rule out the possibility of side effects caused by long-term culture, we isolated MEFs from *Mfn2^flox/flox^* mice. The addition of Cre-expressing adenovirus induced loss of MFN2 within 48 hours, reproduced the rounded morphology, and altered actin cytoskeleton organization seen in *Mfn2*-null MEFs (Fig. S1B-D).

**Figure 1.**
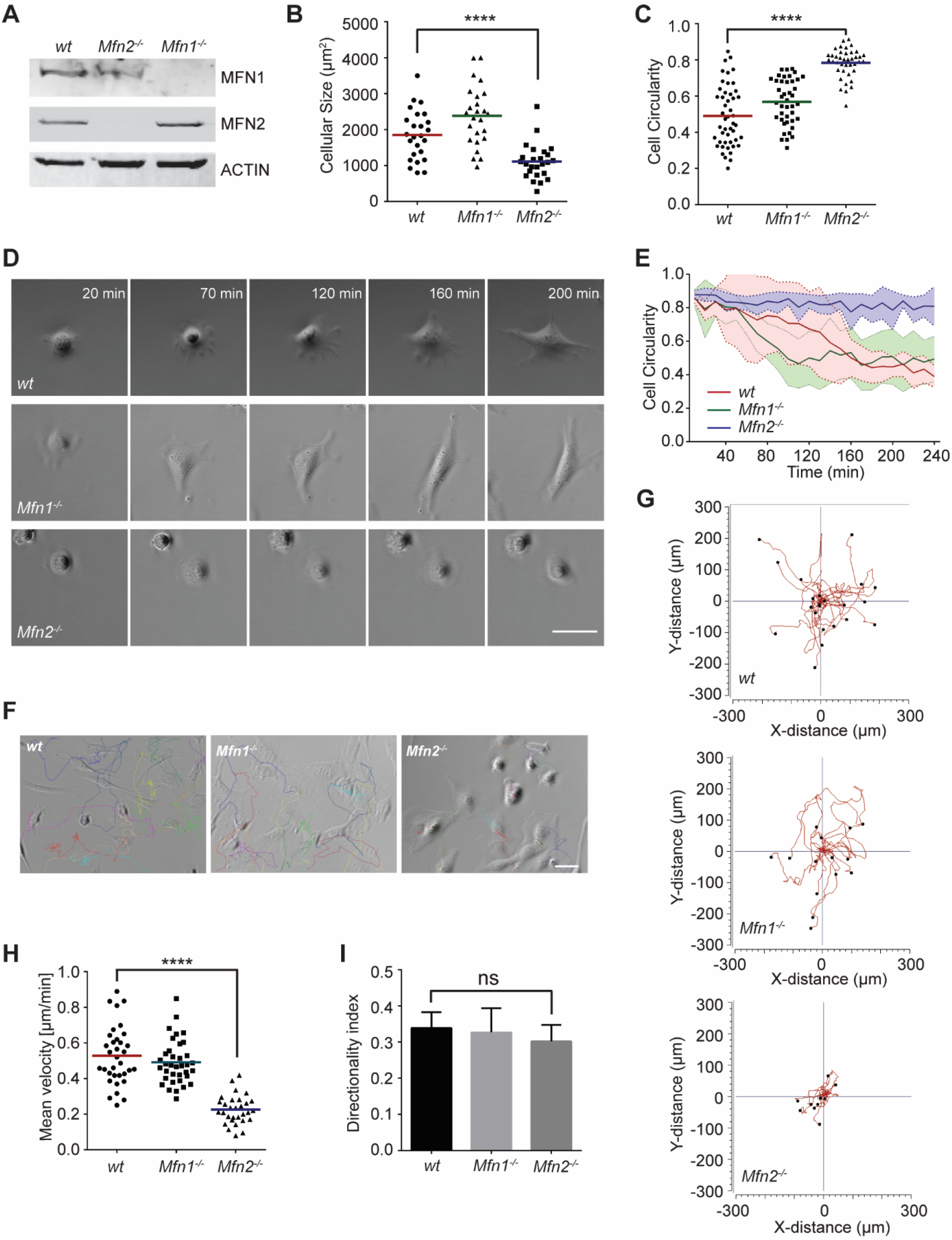
MFN2 regulates cell random migration and cell spreading in MEFs. (A) Western blot determining the expression levels of MFN1 and MFN2 in *wt, Mfn2*-null, and *Mfn1*-null MEFs. (B, C) Cell size (B) and circularity (C) of *wt*, *Mfn1* -null, and *Mfn2*-null MEFs measured after overnight culture. The individual points stand for the size or circularity of individual MEF cells. (D, E) Quantification of cell circularity (E) and representative images (D) of indicated MEFs during cell spreading at indicated time points. Data are presented as mean ± SD in E and were pooled from a total of 6 cells in three independent experiments. (F-H) Representative images with individual tracks (F) and quantification of velocity (H) of MEF cells during random migration. The individual points are mean speeds for individual MEF cells. (G) Wind-Rose plots depicting migratory tracks of *wt*, *Mfn1*-null, and *Mfn2*-null MEFs. (I) Consecutive gene disruption of *Mfn1* or *Mfn2* does not affect directionality. Bars represent arithmetic means ± SD. One representative result of three biological repeats is shown in A, D, F and G. Data are pooled from three independent experiments in B-C, H-I. n>30 cells are tracked and counted in C and D. n>25 cells are quantified in B and H. ****P<0.0001 (one-way ANOVA). Scale bars: 50 μm.

To further confirm the functional role of the MFN2 on cytoskeletal organization and cell migration, we re-expressed MFN1 or MFN2 in *Mfn2*-null MEFs (Fig. 2A). Only MFN2 reexpression significantly increased cell motility (0.32 ± 0.18 μm/min), comparing to *Mfn2*-null MEFs (0.22 ± 0.18 μm/min), or those with MFN1 reexpression (0.21 ± 0.15 μm/min) (Fig. 2B-D, Movie S2). Notably, MFN2 reexpression also rescued the cell’s ability to polarize during the spreading process (Fig. 2E), increased cell area, and decreased circularity (Fig. 2F-G). Similarly, doxycycline (DOX) induced reexpression of MFN2 for 48 hours in *Mfn2*-null MEFs also significantly restored the actin cytoskeleton organization and cell morphology (Fig. 2H-J). Together, these results suggest that these cells’ morphological reorganizations and migratory defects are specifically caused by the loss of MFN2 protein in MEFs.

**Figure 2.**
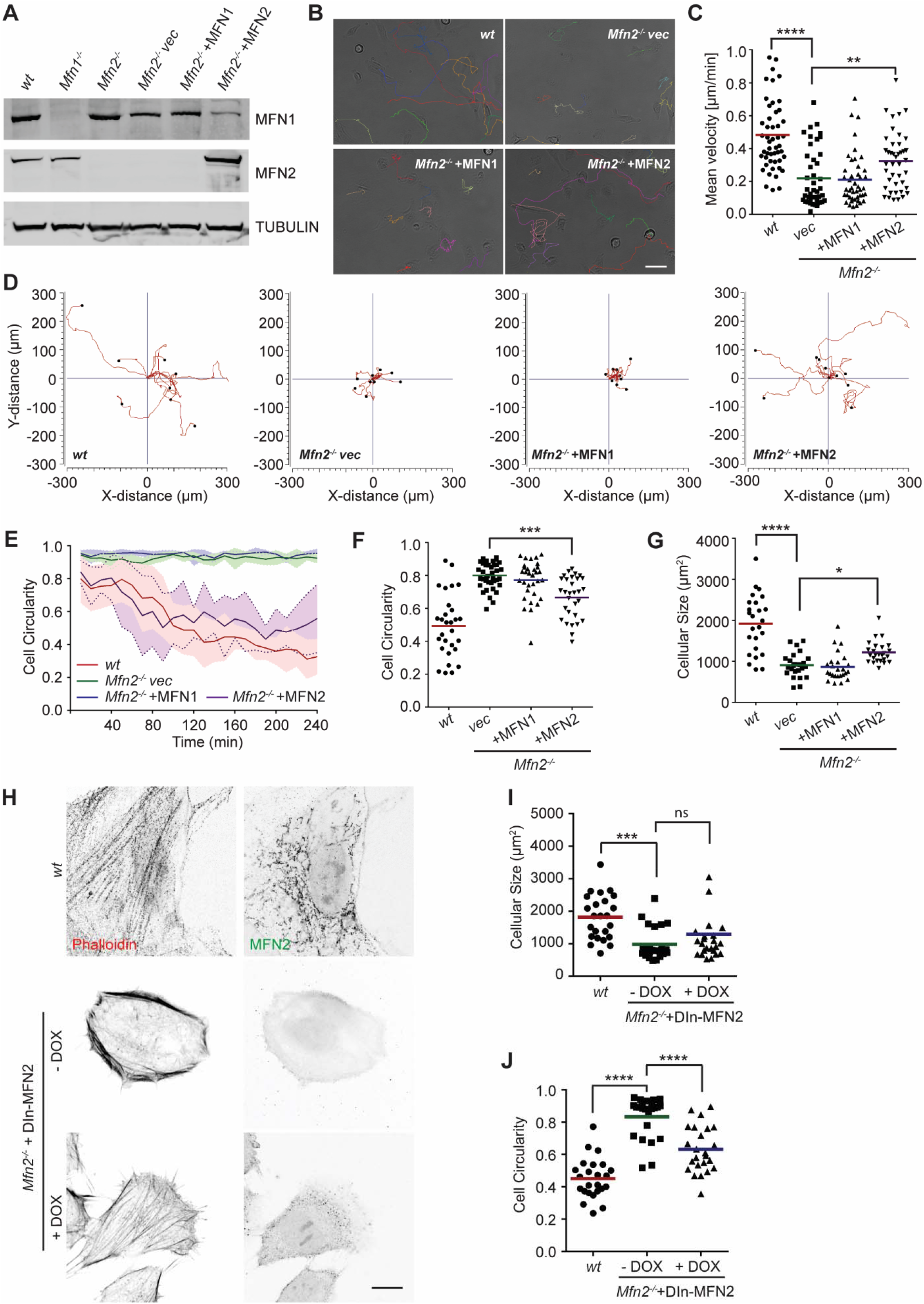
Re-expression of MFN2 rescues cell random migration and cell spreading in *Mfn2*-null MEFs. (A) Western blot determining the expression level of MFN1 and MFN2 in indicated MEF cells. (B, C) Representative images with individual tracks (B) and quantification of velocity (C) of indicated MEF cells during random migration. The individual points are mean speeds for individual MEF cells. (D) Wind-Rose plots depicting migratory tracks of each cell groups in B. (E) Quantification of *wt* and *Mfn2*-null MEFs with vec, MFN1, or MFN2 reexpressed cell circularity during spreading at indicated time points. Data are presented as mean±s.d in E. (n=5). (F, G) Cell circularity (F) and size (G) of indicated MEFs measured after overnight culture. (H-J) Representative images (H) of *wt*, *Mfn2*-null with DIn-MFN2 MEF cells treated with or without doxycycline for 48 hours. The cells are immunostained with phalloidin and MFN2. (I, J) Cell size (I) and circularity (J) of indicated cells in H. The individual points stand for the circularity or size of individual MEF cells in F-G, I-J. One representative result of three biological repeats is shown in A, B, D and H. Data are pooled from three independent experiments in C, F-G, I-J. n>30 cells are tracked and counted in C and F; n>25 cells are quantified in G, I-J. *\*P≥0.05, **P≥0.01*, \*\*\**P*≤0.001, ****P<0.0001 (one-way ANOVA in C, E-F, unpaired t-test in H-I). Scale bars: 50μm in B, 10μm in H.

The significant differences in actin architecture in *Mfn2*-null MEFs, an atypical “Peripheral Actin Band” (PAB), could account for the spreading and migratory defects (Fig. S1A-B, Fig. 2H).

### Increased Cytosolic Ca^2+^ suppresses cell migration in *Mfn2*-null MEFs

These striking alternations in cell morphology, spreading, and migration prompted us to investigate the downstream effectors of MFN2. We focused on one of the MFN2-specific functions: maintaining cellular Ca^2+^ homeostasis by tethering ER and mitochondria. It was previously reported that the increased distance between ER and mitochondria in the absence of MFN2 elevates cytosolic Ca^2+^ transients in MEFs (de Brito and Scorrano, 2008b; Filadi et al., 2015b; Naon et al., 2016b). We, therefore, measured cytosol Ca^2+^ in response to PDGF-BB stimulation and confirmed the previous observation (Fig. 3A). To evaluate the effect of cytosolic Ca^2+^ accumulation on MEF cell migration, we treated *wt* MEFs with the calcium ionophore A23187, an ion carrier facilitating Ca^2+^ transport across the plasma membrane. A reduction of cell migration speed was observed in a dose-dependent manner, which phenocopied the motility reduction in *Mfn2*-null MEFs (Fig. 3B). On the other hand, the reduced motility in *Mfn2*-null MEFs was partially rescued when treated with an intracellular calcium chelator BAPTA-AM (Fig. 3C-D, Movie S3).

**Figure 3.**
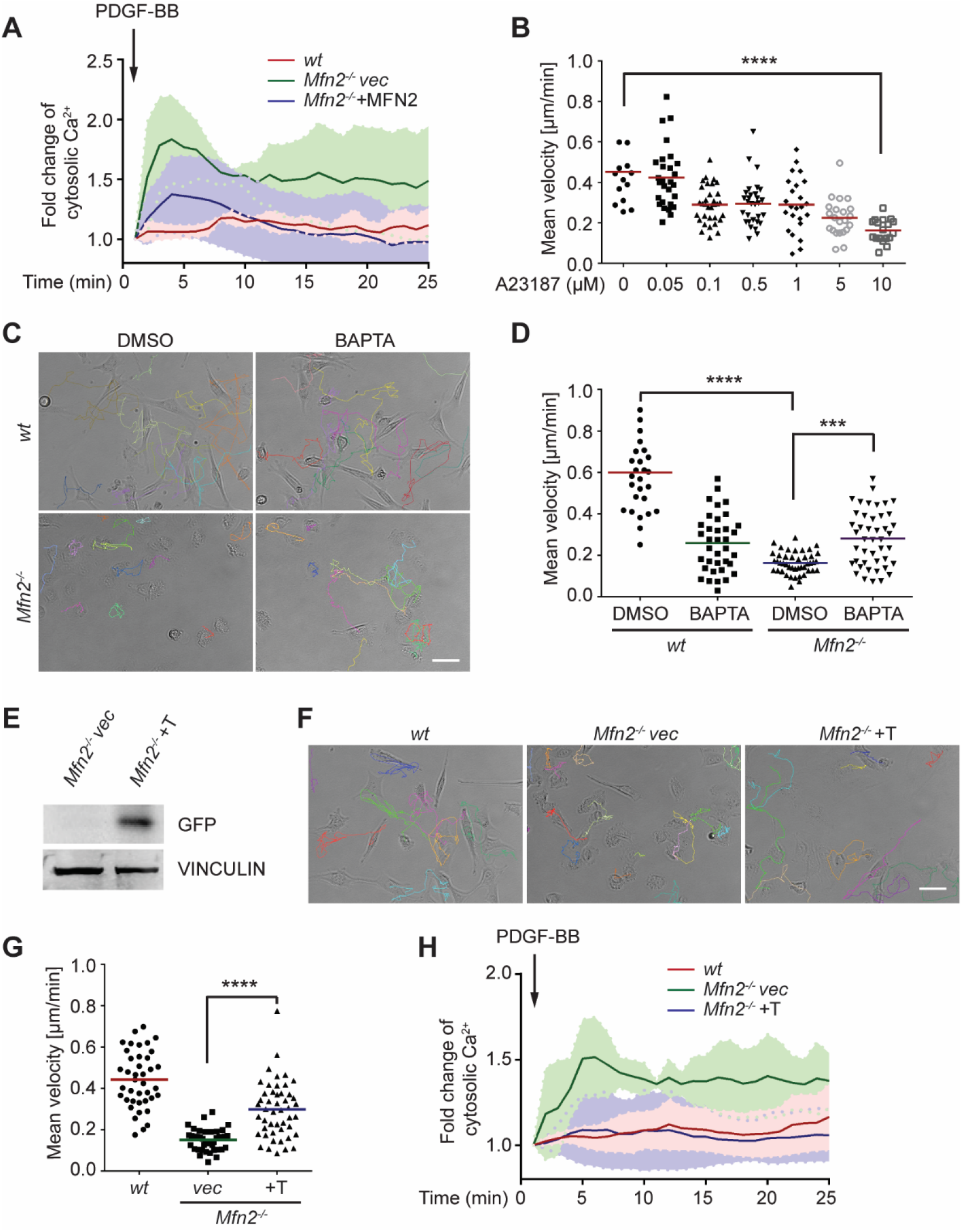
MFN2 regulates random cell migration through cytosolic Ca^2+^ and ER-mitochondria tethering. (A) Fluo-4 recordings of cytosolic Ca^2+^ in the indicated cell lines after PDGF-BB stimulation. (B) Quantification of the velocity of *wt* MEF random migration in the presence of vehicle or different concentrations of the Ca^2+^ ionophore A23187. (C, D) Representative images with individual tracks (C) and quantification of velocity (D) of *wt* or *Mfn2*-null MEFs during random migration with or without the presence of the intracellular calcium chelator BAPTA-AM. (E) Western blot of GFP in indicated cell lines. *Mfn2^-/-^+T, Mfn2*-null MEFs with synthetic ER-GFP-mitochondria tether construct. (F, G) Representative images with individual tracks (F) and quantification of velocity (G) of indicated MEF cells. (H) Fluo-4 recordings of cytosolic Ca^2+^ in the indicated cell lines after PDGF-BB stimulation. The individual points in B, D, and G are mean speeds for individual MEF cells. Data are presented as mean ± SD in A, and H. Data are pooled from three independent experiments in A and H. One representative result of three biological repeats is shown in C-G. n>12 cells are tracked and counted in B. n>25 cells are tracked and measured in D and G. ***P<0.001, ****P<0.0001 [oneway ANOVA (B, G), and two-way ANOVA (D)]. Scale bars: 50 μm.

To confirm that MEF2 regulates cytosolic Ca^2+^ via maintaining ER-mitochondria tether, we introduced an artificial tether construct (Kornmann et al., 2009) into the *Mfn2*-null MEFs. The tether comprises a GFP protein carrying ER and mitochondrial localization sequences at opposite ends, which functions independently of MFN2. Expressing the tether corrected the cytosolic Ca^2+^ levels in response to PDGF-BB stimulation and partially rescued the migration speed and cell morphology in *Mfn2*-null MEFs (Fig. 3F-H, Movie S4). These data suggest that MFN2 regulates cell morphology and adhesive migration in MEFs by maintaining mitochondria-ER interaction and Ca^2+^ homeostasis.

### Elevated CaMKII activation is associated with MFN2-regulated random migration

Given that cytosolic Ca^2+^ plays essential roles in MFN2-mediated cytoskeleton regulation and cell migration, we looked at the kinases and phosphatases regulated by Ca^2+^/ calmodulin (CaM), including CaMKK, CaMKII, and Calcineurin, which are previously shown to regulate actin bundling (Saneyoshi and Hayashi, 2012) (Fig. 4A). AIP, the CaMKII inhibitor, partially restored motility in *Mfn2*-null MEFs. Neither the calcineurin inhibitor FK506 nor the CaMKK inhibitor STO609 had any effect (Fig. 4B-C). We found a higher level of phosphorylated or active CaMKII in *Mfn2*-null MEFs, which was also reduced by the expression of MFN2 or the tether (Fig. 4D-E). The function of CamKII was further confirmed by expressing wild-type (CaMKII-WT) or dominant negative (CaMKII-DN) versions of CamKII in *Mfn2*-null MEFs. CaMKII-DN, but not CaMKII-WT, induced a moderate but significant increase in migration velocity (Fig. 4F-G). Together, our results indicate a role of CaMKII in cytoskeleton regulation in *Mfn2*-null MEFs.

**Figure 4.**
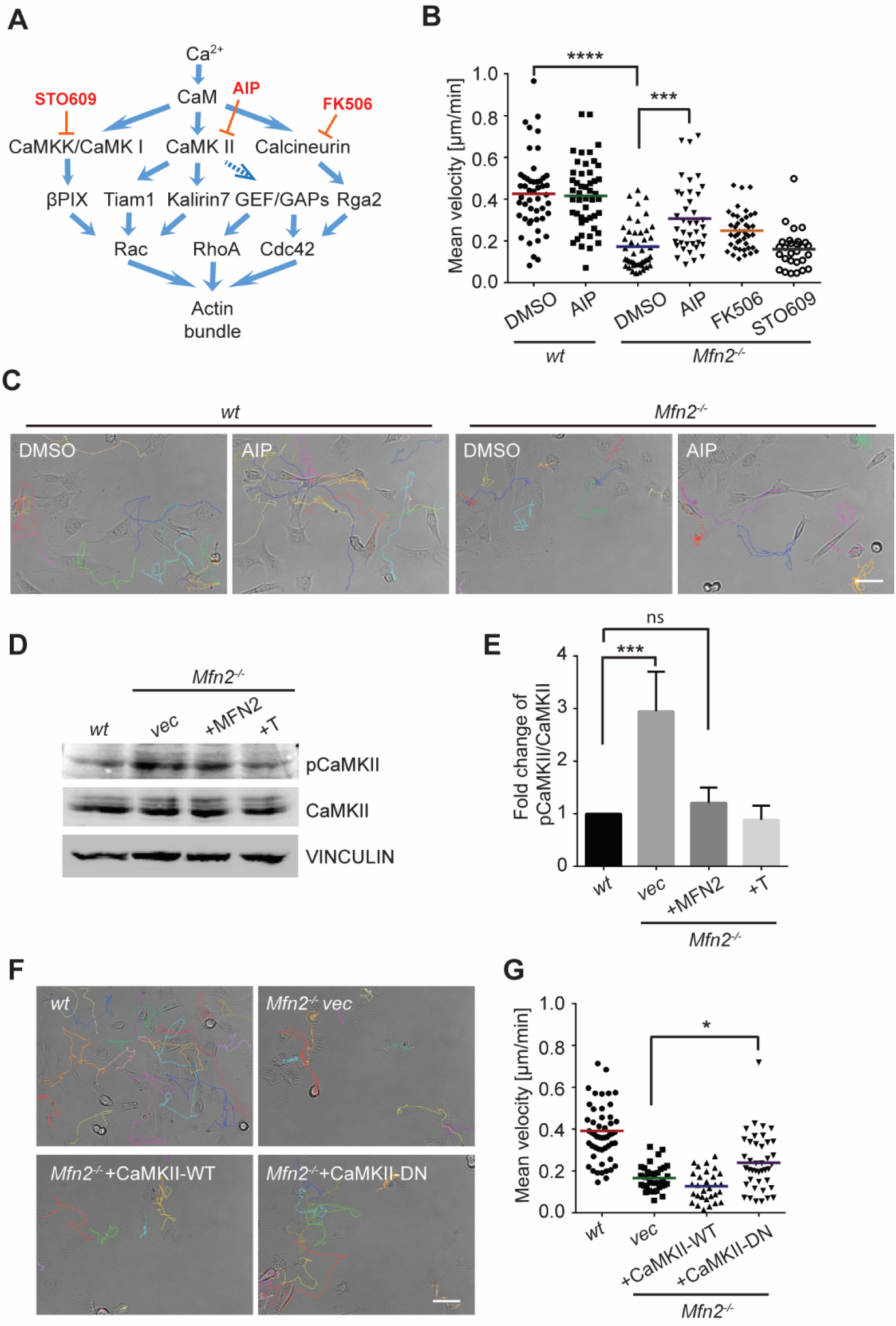
CaMKII activation mediates Mfn2 deficiency-induced inhibition in MEF migration. (A) Selected signaling cascades involved in the regulation of the actin cytoskeleton via Ca^2+^. Blue arrows indicate positive regulation. Dashed blue arrows indicate positive regulation with unclear mechanisms. Orange T-shape bars indicate negative regulation of the pharmacological inhibitors. (B) Quantification of the velocity of indicated MEF cells treated with DMSO, the CaMKII inhibitor AIP, the Calcineurin inhibitor FK506, or the CaMKK inhibitor STO609. (C) Representative images with individual tracks of *wt* or *Mfn2*-null MEFs treated with the CaMKII inhibitor AIP. (D, E) Western blot (E) and quantification (F) determining the amount of pCaMKII and pan-CaMKII in *wt*, *Mfn2*-null MEFs with vec, CaMKII-WT, or CaMKII-DN overexpressed after treating with 25 μM PDGF-BB for 4 mins. (F, G) Quantification of velocity (G) and representative images with individual tracks (F) of *wt, Mfn2*-null MEFs with vec, CaMKII-WT, or CaMKII-DN overexpressed during random migration. One representative result of three biological repeats is shown in C-G. n>30 cells are quantified in B and G. *P≤0.05, ***P≤0.001, ****P<0.0001[one-way ANOVA (E, G), and twoway ANOVA (B)]. Scale bars: 50 μm.

### MFN2 deficiency-induced migration defect is independent of Rac and CDC42

The Rho GTPase family members are master regulators of the actin cytoskeleton and cell migration. The peripheral enrichment of actin filaments and extensive membrane ruffles we observed in *Mfn2*-null MEF cells resembled the classic phenotype seen in fibroblasts with constitutively active Rac(Hall, 1998a). Therefore, we hypothesized that Rac might be overactivated in MFN2-depleted MEF cells. We performed a RAC-GTP pulldown and did not observe any significant increase in the absence of MFN2 (Fig. S2A-B). In addition, the Rac inhibitor, CAS1090893, did not rescue the cell motility defect of the *Mfn2*-null MEFs (Fig. S2C), suggesting that Rac is not the primary effector regulated by MFN2. CDC42-GTP levels are comparable between the *wt* and *Mfn2*-null cells (Fig. S2D-E).

### Loss of MFN2 drives overactivation of RhoA GTPase and redistribution of focal adhesions

In contrast to the minor changes in Rac and CDC42 activity, we detected a marked increase of RhoA-GTP in *Mfn2*-null MEFs (Fig. 5A-B). Restoring MFN2 expression or introducing the ER-mitochondria artificial tether brought the RhoA activation level back to the *wt* level. We then compared the distribution of focal adhesion protein paxillin (Pax) by immunofluorescence. In addition to the striking PAB architecture described previously, the focal adhesion complexes in *Mfn2*-null MEFs appeared larger, consistent with the observation of RhoA overactivation in fibroblasts (Hall, 1998b). The focal adhesions were also restricted to the cell periphery in *Mfn2*-null MEFs (Fig. 5C). We plated the cells on fibronectin, fibrinogen, or uncoated cover glasses and discovered that the PAB formation is independent of the extracellular substrates (Fig. S3A).

**Figure 5.**
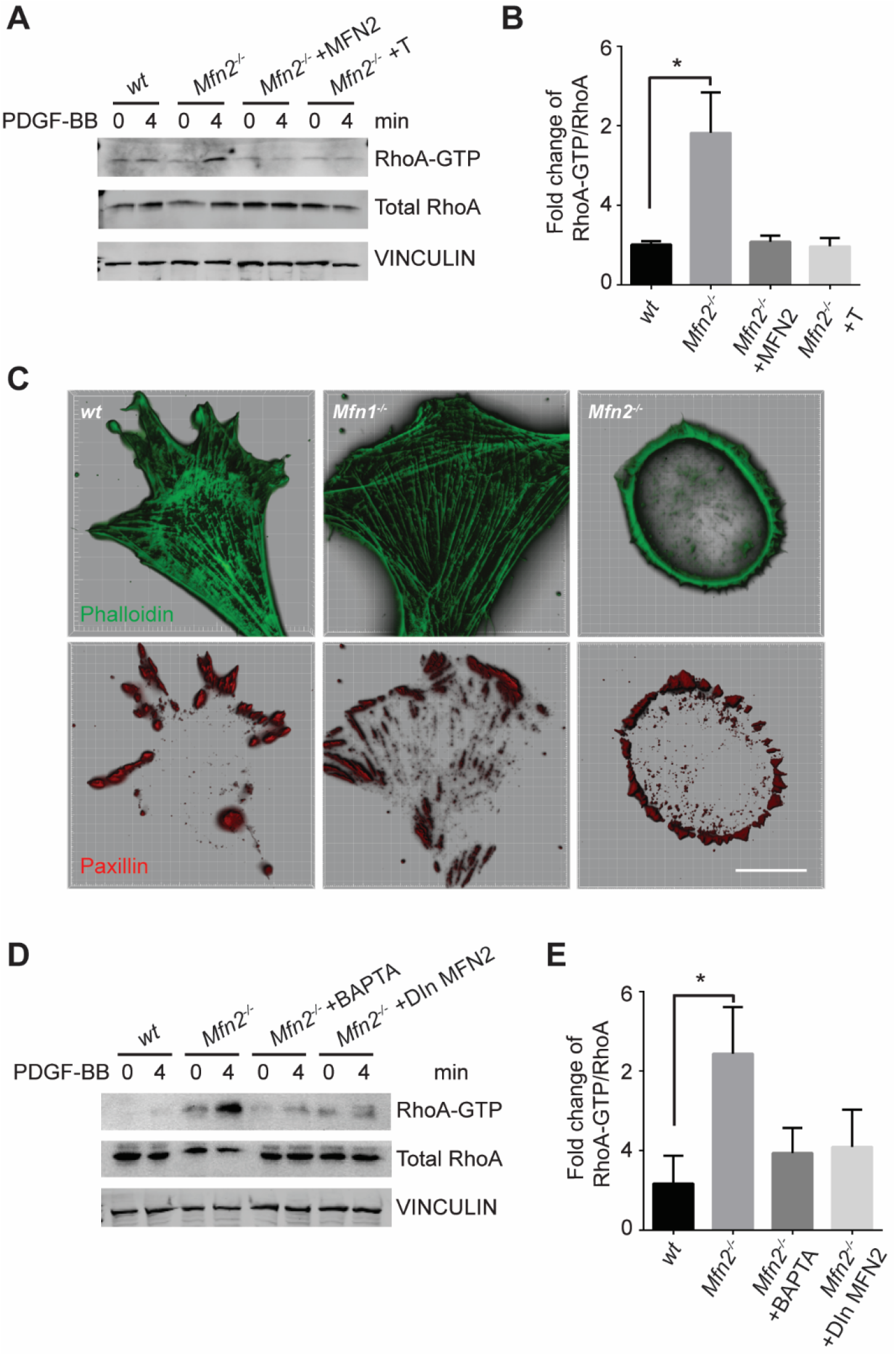
Loss of MFN2 induces heightened RhoA activation in MEFs. (A, B) RhoA pulldown activation assay demonstrates increased RhoA-GTP in *Mfn2*-null MEFs, which can be corrected by re-expressing MFN2, inducing a mitochondria-ER tether. (A) Western blot and (B) quantification determining the amount of RhoA-GTP and total RhoA protein in *wt*, *Mfn2*-null MEFs, *Mfn2*-null MEFs with MFN2 re-expression, or with an artificial ER-mitochondria tether; the indicated cell lines were treated with 25 ng/ml PDGF-BB for 0 or 4 minutes. (C) Immunofluorescence of F-actin (phalloidin) and Paxillin in *wt*, *Mfn1*-null, and *Mfn2*-null MEFs after overnight culture. (D, E) RhoA pulldown activation assay demonstrates cytosolic Ca^2+^ inhibition corrects RhoA-GTP level *in Mfn2*-null MEFs. (D) Western blot and (E) quantification showing the amount of RhoA-GTP protein in *wt*, *Mfn2*-null MEFs, *Mfn2*-null MEFs treated with BAPTA, or *Mfn2*-null MEFs with DOX-induced MFN2 re-expression for 48 hours; the indicated cells were treated with 25ng/ml PDGF-BB for the indicated time. One representative result of two biological repeats is shown in A, C, and D. \**P*≤0.05 (one-way ANOVA comparing each group with the average of *wt* group). Scale bar: 50 μm.

These focal adhesion differences hinted that the spreading and migration defects in cells depleted of MFN2 might be explained, at least in part, by significantly increased RhoA activity. Moreover, BAPTA-AM treatment or MFN2-reexpression reduced the heightened activity of RhoA in *Mfn2*-null MEFs (Fig. 5D-E), indicating that the cytosolic Ca^2+^ increase was responsible for the RhoA overactivation in *Mfn2*-null MEFs.

### Increased Myosin Regulatory Light Chain activity is critical for PAB formation in *Mfn2-* null MEFs

To fully understand the downstream mechanism of MFN2 in regulating the cytoskeleton, we used pharmacological inhibitors for proteins regulated by CaM (Fig. 6A, Fig. S3B,C). The RhoA inhibitor I, the Rho-associated protein kinase (ROCK) inhibitor Y27632, the myosin inhibitor Blebbstatin, and the myosin light chain kinase (MLCK) inhibitor ML-7 showed the most pronounced effects on restoring both the motility and cell morphology in *Mfn2*-null MEFs (Fig. 6A-B, Fig. S3B-C, Movie S5). The focal adhesion kinase inhibitor 14 (FAK inhibitor 14), the LIM kinase (LIMK) inhibitor BMS-5, and the Arp2/3 inhibitor CK-666 had no statistically significant effects on restoring cell motility (Fig. S3B). This result was confirmed by analyzing the actin filament organization and focal adhesion distribution in inhibitor-treated *Mfn2*-null MEFs. RhoA inhibitor-I, ML-7, Y27632, or Blebbstatin abrogated the PAB *in Mfn2*-null MEFs and restored typical fibroblast characteristics, including the formation of filopodia, developed cell edges, and focal adhesions at the leading and trailing edges (Fig. 6C). Taken together, these findings demonstrate a potential mechanism whereby excessive cytosolic Ca^2+^ in the absence of MFN2 leads to the overactivation of RhoA and MLCK, which increases MLC activity and contributes to the PAB in *Mfn2*-null MEFs (Fig. 6D, Fig. S3D).

**Figure 6.**
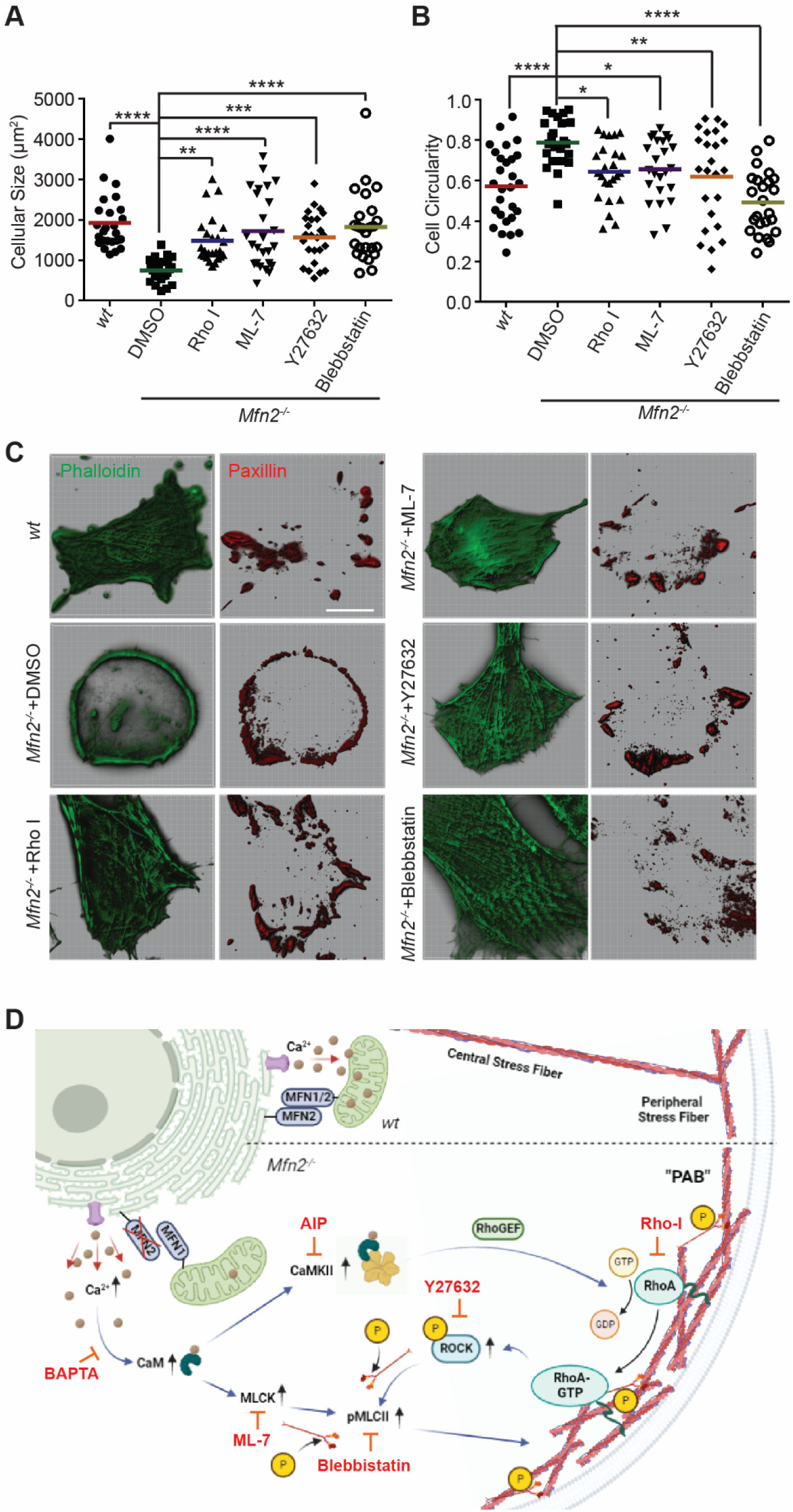
Small molecule inhibitors targeting RhoA- and MLCK-related signaling pathways rescue MFN2 deficiency-induced phenotypes. (A, B) Cell size (A) and circularity (B) of *wt* and *Mfn2*-null MEFs treated with indicated inhibitors overnight. The individual points stand for the size or circularity of individual MEF cells. (C) Immunofluorescence of F-actin (phalloidin) and Paxillin in *wt* and *Mfn2*-null MEFs treated with indicated inhibitors overnight. (D) Schematic of the effectors and their inhibitors (red) in MFN2 regulated signaling network leading to Actin bundle. Increased cytosolic Ca^2+^ may activate MLCK, CaMKII, and RhoA-ROCK, which activate MLC and affect actin bundle formation. One representative result of three biological repeats is shown in C. Data are pooled from three independent experiments, and *n*>30 cells are quantified in A and B.\**P*≤0.05, \*\**P*≤0.01, \*\*\**P*≤0.001, \*\*\*\**P*<0.0001 (one-way ANOVA comparing each group to the average of *Mfn2*^-/-^ DMSO group). Scale bars: 50 μm.

Given the essential role of the MLC, we probed for pMLCII in *wt*, *Mfn1* -null, and *Mfn2*-null MEFs. As expected, the pMLCII level was significantly increased in *Mfn2*-null MEFs (Fig. 7A-B), which could be corrected by expressing MFN2 or the artificial tether in *Mfn2*-null MEFs (Fig. 7C-D). Using immunofluorescence microscopy, we noted that pMLCII colocalized with peripheral actin bundles in *Mfn2*-null MEFs (Fig. 7E). To confirm the importance of MLC activity, we knocked down MLCK or ROCK in *Mfn2*-null MEFs (Fig. 7F). Both knockdown cell lines displayed significantly reduced pMLCII levels (Fig. 7G-H), restored stress fiber architecture, redistributed focal adhesions, and restored protrusive structures such as filopodia (Fig. 7I). Both knockdown lines also showed an increase in cell size and greater polarization compared to *Mfn2*-null cells (Fig. 7J-K).

**Figure 7.**
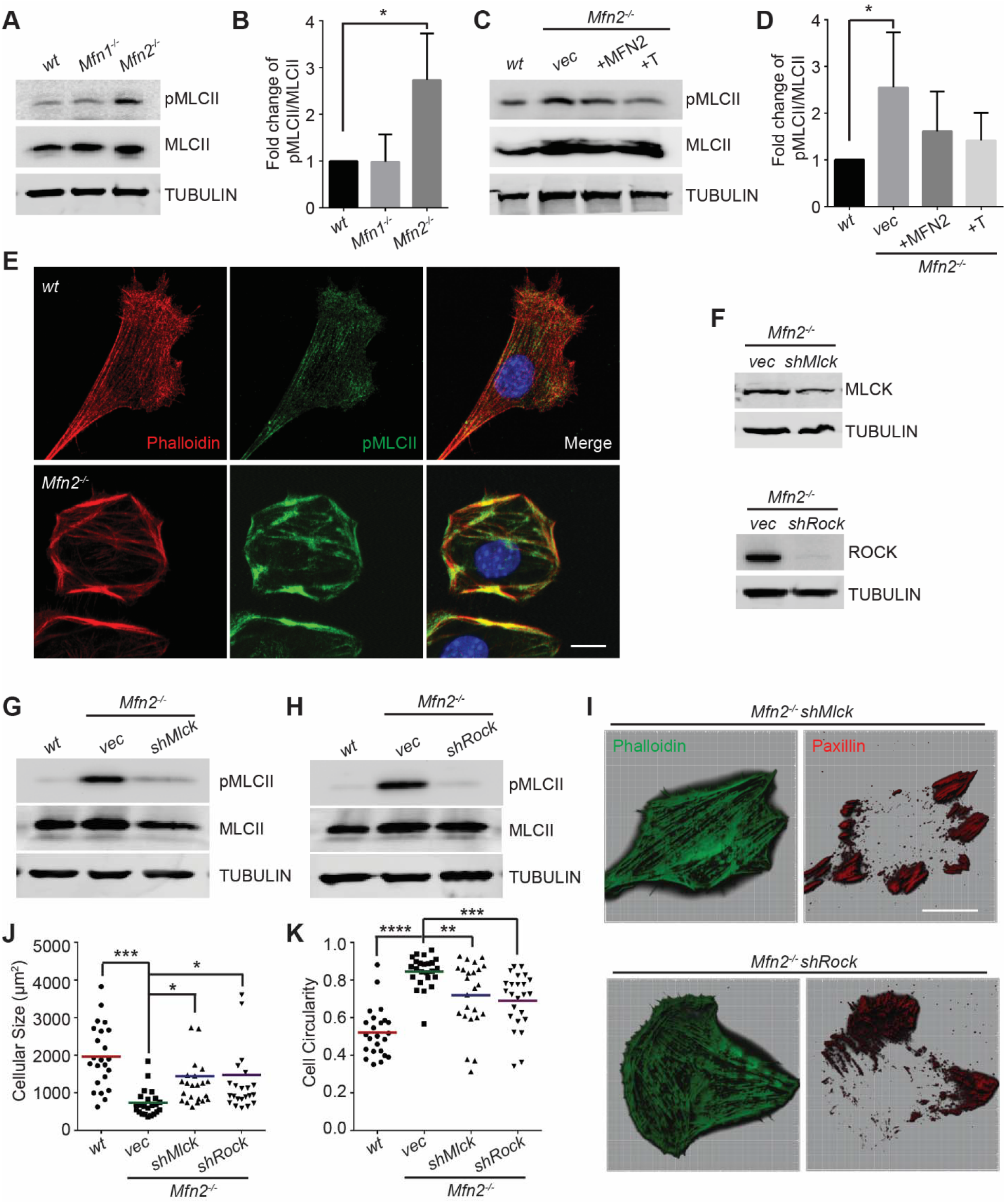
Heightened MLC activity promotes the “PAB” structure in MFN2 deficient MEFs. (A, B) Western blot (A) and quantification (B) of the amount of pMLCII and total MLCII in *wt*, *Mfn1* -null, and *Mfn2*-null MEFs. (C, D) Increased pMLCII in *Mfn2*-null MEFs can be corrected by re-expressing MFN2 or inducing a mitochondria-ER tether. (C) Western blot and (D) quantification determining the amount of pMLCII and total MLCII protein in *wt*, *Mfn2*-null MEFs, *Mfn2*-null MEFs with MFN2 re-expressed, or with an artificial ER-mitochondria tether. (E) Representative images of *wt* and *Mfn2*-null MEFs immunostained for F-actin (phalloidin), pMLCII, and DAPI. (F) Western blot determining the expression levels of MLCK or ROCK in *Mfn2*-null MEFs with *shMLCK* or *shROCK*. (G, H) Western blot of pMLCII and total MLCII *Mfn2*-null MEFs with *shMLCK* or *shROCK*. (I) Representative images of *Mfn2*-null MEFs with *shMLCK* or *shROCK* immunostained for F-actin (green) and paxillin (red). (J, K) Cell size and circularity of *wt*, *Mfn2*-null MEFs with *vec, shMLCK*, or *shROCK* were measured after overnight culture. The individual points stand for the size or circularity of individual MEF cells. One representative result of three biological repeats is shown in A-I. Data are pooled from three independent experiments in J-K. *n*>25 cells are tracked and counted in J and K. \**P*≤0.05, \*\**P*≤0.01, \*\*\**P*≤0.001, \*\*\*\**P*<0.0001 (one-way ANOVA, comparing each group to the average of *Mfn2^-/-^ vec* group in J-K). Scale bars: 50 μm in I, 10μm in E.

As MFN2 is a mitochondrial protein, we also performed a seahorse assay to measure mitochondrial functions in each cell line. *Mfn2*-null MEFs showed a decreased rate of oxidative metabolism relative to *wt* and *Mfn1*-null MEFs (Fig. S4A, B), while MFN2 reexpression or introducing ER-mitochondrial tether increased the overall oxygen consumption rate in *Mfn2*-null MEFs (Fig. S4C, D). Notably, knocking down MLCK in *Mfn2*-null MEFs enhanced oxidative metabolism, whereas ROCK knockdown further reduced oxygen consumption (Fig. S4E, F). Given that both MLCK and ROCK knockdown can restore cell spreading and migration in *Mfn2*-null MEFs, it is unlikely that alterations in mitochondrial metabolism are the primary determinant in PAB formation.

### Myosin Regulatory Light Chain overexpression phenocopied MFN2 depletion

Mitochondria and MFN2 regulate multiple cellular signaling pathways in addition to cytosolic calcium. To determine whether the PAB is primarily driven by RhoA or MLCK activation (downstream of CaM), we attempted to constitute the PAB in *wt* MEFs. We first tried Rho activator treatment or expression of constitutively active MLCK (MLCK-CA)(Wong et al., 2015) (Fig. 8A). *Wt* MEFs treated with Rho Activator exhibited more and thicker bundles of actin filaments, with a mesh-like structure, across the dorsal side of the cells, but maintained their overall polarized morphology. This observation is consistent with the known function of RhoA to induce the central stress fibers formation in fibroblasts (Katoh et al., 2001b; a). In contrast, MLCK-CA expression increased peripheral stress fibers and noticeably fewer bundles of actin filaments in the central portion but maintained the classic polarized morphology. When combined (Rho activator treatment + MLCK-CA expression), the effect was increased, with the cells’ thick actin bundles along the periphery. These cells, however, retained their protrusive structures and polarized shape (Fig. 8A).

**Figure 8.**
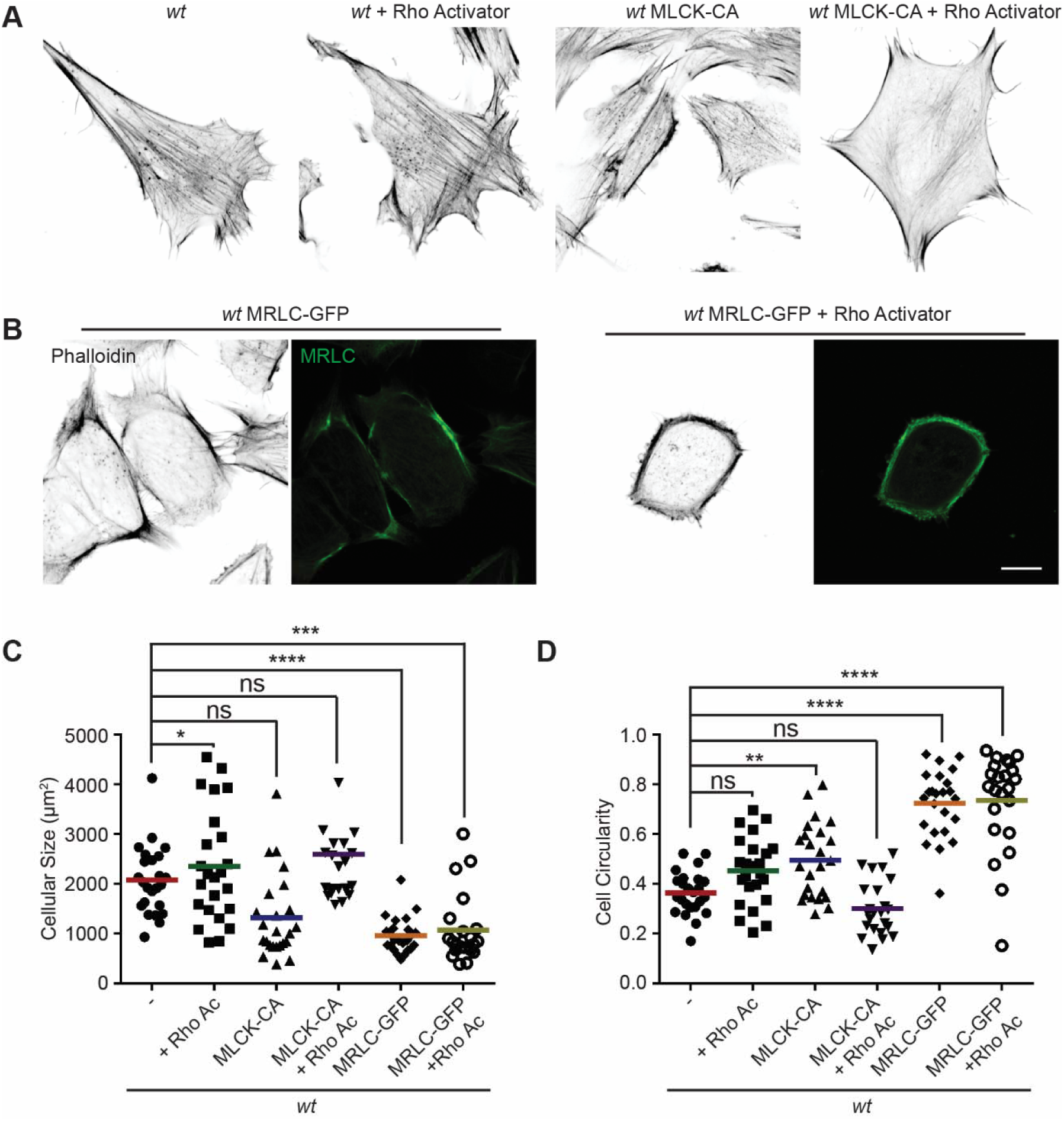
MRLC overexpression and Rho Activation in *wt* MEFs recapitulates the “PAB” structure. (A) Immunofluorescence of F-actin (phalloidin) in *wt* MEFs with or Rho Activator treatment or introducing MLCK-CA expression. (B) Representative images of *wt* MEFs expressing MRLC-GFP with or without the Rho Activator, immunostained for F-actin (phalloidin). (C, D) Cell size and circularity of *wt* MEFs with the indicated overexpression or drug treatment. The individual points stand for the size or circularity of individual MEF cells. One representative result of three biological repeats is shown in A-B. Data are pooled from three independent experiments in C-D. *n*>20 cells are tracked and counted in C and D. \**P*≤0.05, *\*\*P*<0.01, \*\*\**P*≤0.001, \*\*\*\**P*<0.0001 (one-way ANOVA, comparing each column with the mean of *wt* group). Scale bars: 10μm.

To our surprise, when we attempted to image the myosin dynamics in *wt* and *Mfn2*-null MEFs, we noticed a large percentage (55%) of *wt* MEFs with round shapes, as well as peripherally enriched actin filaments after overexpressing the GFP-tagged myosin regulatory light chain (MRLC-GFP) (Fig. 8B). Cells expressing MRLC-GFP were significantly smaller in size and became round in shape (Fig. 8C-D), however still preserved their filopodia and other cell protrusions. When these cells were treated with the Rho activator, a noticeable percentage (48%) displayed the PAB morphology, including a decrease in cell area and an increase in cell roundness (Fig. 8B-D). In summary, the combination of MRLC overexpression with pharmacological activation of RhoA is sufficient to drive PAB formation in MEFs.

### Loss of MFN2 displays different cytoskeletal architecture with increased cell contractile forces

To better virtualize the individual actin filaments in the PAB, we performed 3D super-resolution imaging of F-actin in *wt* and *Mfn2*-null MEFs (Fig. 9A). In *Mfn2*-null MEFs, F-actin are in parallel directions of the cell cortex, in sharp contrast to those in *wt* MEF, protruding against the membrane. We then used atomic force microscopy (AFM) to measure cell stiffness*. Mfn2* -null MEFs showed a softer Young’s modulus than *wt* cells (Fig. S5B), consistent with the previous report that cells with apical stress fibers like fibroblasts are stiffer than cells without (Efremov et al., 2019). We then measured plasma membrane tension using the Flipper-TR dye and FLIM imaging (Colom et al., 2018). Consistently, *Mfn2*-null MEFs showed a lower fluorescence lifetime, indicating lower membrane tension than the *wt* (Fig. S5C). The reduced membrane stiffness and tension are consistent with a less spread cell morphology.

**Figure 9.**
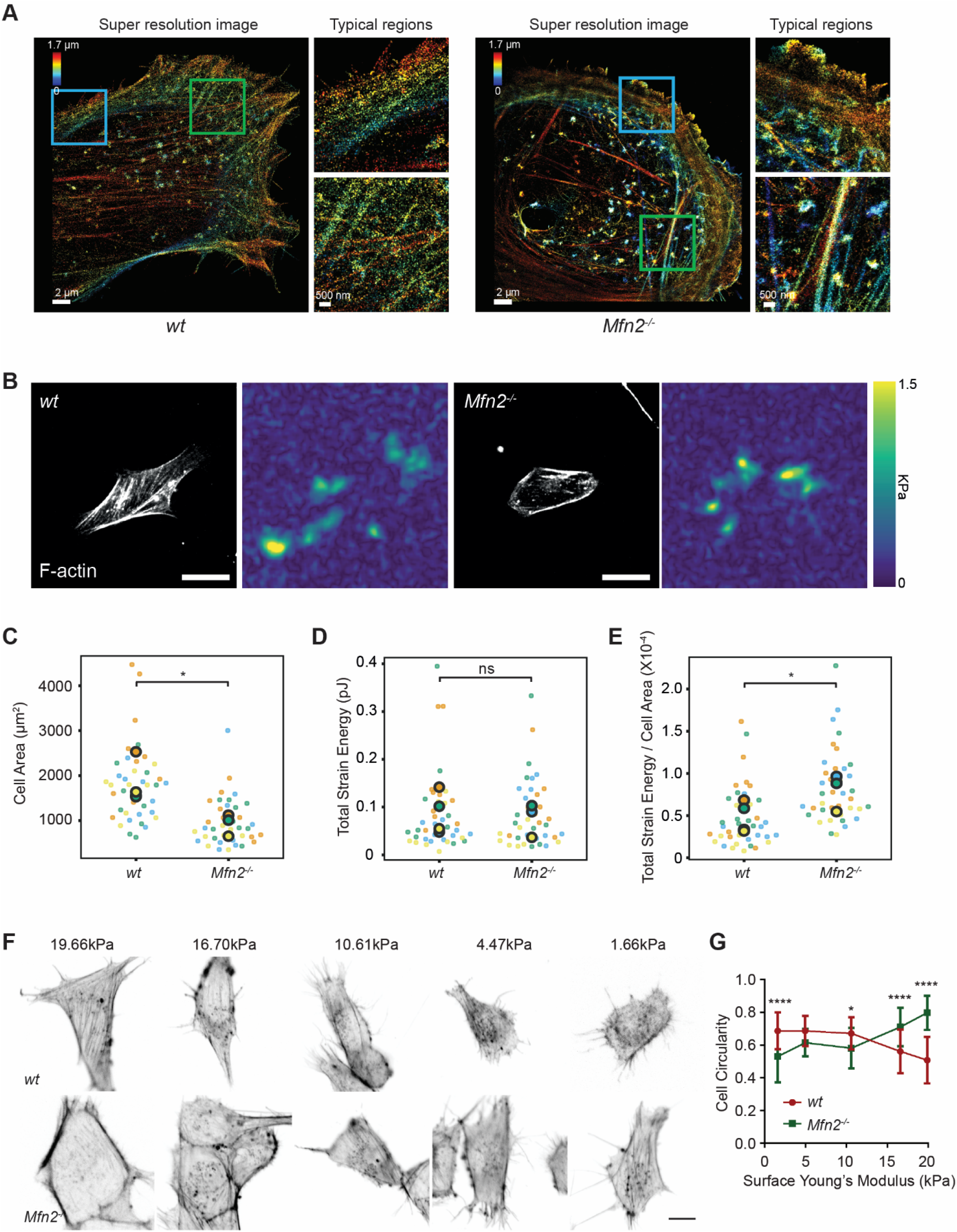
Mfn2-null MEFs exhibit altered actin organization and cell stiffness. (A) 3D super-resolution reconstructions of immunofluorescence-labeled F-actin in *wt* or *Mfn2*-null MEFs. x–y overview of a 1.7-μm-thick volume of the cells. (B) Morphology of indicated MEF cells on polyacrylamide (PAA) substrates after overnight culture immunostained with Alexa-488 phalloidin. A traction stress map with color values corresponding to different stress values is shown on the right. Scale bars: 50 μm. Quantification of the corresponding cell spreading areas (C), total strain energy (D), and total strain energy normalized to cell area (E). (F) Representative images of indicated MEF cells on polyacrylamide (PAA) substrates of different stiffness after overnight culture. The cells are immunostained with Alexa-488 phalloidin. (G) The cell circularity of the indicated cells is measured. The individual points stand for the circularity of individual MEF cells. One representative result of three biological repeats is shown in C and F. Data are pooled from three independent experiments in C-E, and G. n>40 cells are counted in BE. n>25 cells are tracked and measured in J and K. *P≡0.05, ****P<0.0001 (unpaired t-test). Scale bars: 2μm in A, 50μm in B, 10μm in F.

The prominent peripheral F-actin bundled with MLC (Fig. 7E) possibly generates a non-polar, global contraction at the cell periphery. We, therefore, utilized traction force microscopy (TFM) to measure the contractile force of the *Mfn2*-null MEFs (Oakes et al., 2014a). After normalizing to the cell area, the average strain energy in *Mfn2*-null MEFs is significantly higher than with *wt* cells, which correlated with the elevated actin-myosin level. Since fibroblasts generate more traction force on stiffer substrates (Lo et al., 2000), we then asked whether or to which extent substrate stiffness affects the PAB structure in *Mfn2*-null MEFs. We cultured the cells on polyacrylamide (PAA) gels with different stiffness and stained F-actin (Fig. 9F). As previously reported, *wt* cells showed diminished cell spreading and rounder morphology on softer substrates. In contrast, *Mfn2*-null MEFs displayed a more elongated cell shape on softer substrates, with partially restored stress fibers at the cell center (Fig. 9F, G). These results suggest that substrate stiffness affects cell spreading, and strong substrate interaction or outside-in signal is required for the “PAB” structure in *Mfn2*-null MEFs.

## Discussion

Our results show that loss of MFN2 protein results in defective spreading and polarization, along with reduced motility of MEF cells in 2D migration. This phenotype results from an increased cytosolic Ca^2+^ level upon Mfn2 removal and loss of ER-mitochondria tethers, which lead to the higher activity of calcium-regulated kinases, including CaMKII and MLCK, overactive RhoA and NMII, and an accumulation of peripherally localized actin and myosin, which we named the PAB. The cell morphology of *Mfn2*-null MEFs can be rescued by restoring MFN2 expression, introducing ER-mitochondria artificial tethers, or inhibiting cytosolic Ca^2+^, CaMKII, MLCK, RhoA, or MRLC. Thus, together, these data identify the mechanism for how Mfn2 regulates cell spreading and adhesive migration in MEFs and highlight its essential function in maintaining mitochondria-ER contact and regulating the actomyosin network.

In line with the “PAB” structure and markedly enriched myosin at the cell periphery in *Mfn2*-null MEFs, they displayed higher strain energy as determined by TFM. The findings suggest that an increase in peripheral traction force hampered *Mfn2*-null MEFs from further spreading. However, this effect could be corrected by reducing substrate stiffness, given that a certain percentage of *Mfn2*-null MEFs gained elongated morphology on soft substrates. It is well known that mechanical forces play a significant role in regulating cell adhesion and cytoskeletal organization (Discher et al., 2005; Burridge and Chrzanowska-Wodnicka, 2003). Cells generate higher traction stresses on stiffer substrates, and contractility-induced tension drives the formation of stress fibers and focal adhesions. Conversely, stress fibers and focal adhesions disassemble when contractility is inhibited (Burridge and Chrzanowska-Wodnicka, 2003; Pelham and Wang, 1997; Wong et al., 2015). A possible explanation of our observation is that the “outside-in” feedback loop coupling with the elasticity of the extracellular microenvironment abrogated the aberrant “PAB” architecture.

Cell migration is a highly dynamic process in which actin treadmilling and focal adhesion turnover orchestrate front-to-rear polarity and cell movements. Rac is known for its regulatory functions in the formation of focal complexes and lamellipodia at the front(Hall, 2005; Nobes and Hall, 1999, 1995), while RhoA modulates actomyosin contraction (Ridley and Hall, 1992) and focal adhesion disassembly (Ridley and Hall, 1992; Nobes and Hall, 1999, 1995) at the rear end. Our previous research proved that MFN2 suppresses Rac activation and supports cell adhesion in neutrophils (Zhou et al., 2021). However, we found that RhoA, instead of Rac, plays a dominant function in actin cytoskeleton reorganization in *Mfn2*-null MEFs. This difference is not entirely surprising, given that these two cells utilize different migratory modes. Neutrophils are a type of fast amoeboid-migrating cell that don’t form mature focal adhesions during cell migration even on specific substrates(Lämmermann et al., 2008; de Bruyn, 1946). In contrast, slower-moving cells, like fibroblasts, form mature focal adhesions and require focal adhesion recycling to move (Lauffenburger and Horwitz, 1996; Seetharaman and Etienne-Manneville, 2020). Despite described controversies concerning the dominant downstream effectors in neutrophils and fibroblasts, the proposed negative regulatory function of MFN2 on cytosolic Ca^2+^ levels and its functional role in cell migration was consistent in both cell lines.

The elevation of cytosolic Ca^2+^ concentration is known to induce the activation of MLCK. MLCK phosphorylates the 20 kDa regulatory myosin light chain (MLC) and consequently activates the myosin ATPase activity (Ikebe and Hartshorne, 1985). The Rho-kinase, ROCK, also controls MLC activity by phosphorylating at Ser19 and Thr 18 (Kassianidou et al., 2017; Totsukawa et al., 2000). It is reported that the biphosphorylated MLC (pp-MLC) localizes to the cell center. In contrast, the monophosphorylated MLC (p-MLC) tends to locate in the cell periphery(Kassianidou et al., 2017). Indeed, this is consistent with our observation that the pp-MLC level did not change when we probed pp-MLC with an antibody specific for both Ser19 and Thr18 (data not shown). Only MLC pSer19 was markedly elevated in *Mfn2*-null MEFs. However, interestingly, either MLCK inhibitor ML-7 or ROCK inhibitor Y27632 restored cell morphology and motility in MEFs without MFN2. Our data suggested that both ROCK and MLCK are required for enhanced MLC phosphorylation at Ser19 in the MFN2-null MEFs.

The RhoA/ROCK signaling pathway plays an essential role in response to cytosolic Ca^2+^ (Uehata et al., 1997; Saneyoshi and Hayashi, 2012; Ying et al., 2009). PDZ-RhoGEF is a vital effector in response to cytosolic Ca^2+^ (Ying et al., 2009; Derewenda et al., 2004). Cytosolic Ca^2+^ activates RhoA through the PYK2/PDZ-RhoGEF pathway in five cell lines (Primary rat aortic vascular smooth muscle cells, HEK293T, MDCK, Neuron2A, and PC12)(Ying et al., 2009). In line with the previous observation(Ying et al., 2009), BAPTA-AM abolishes RhoA activation in our current work. Further work will be required to determine whether PDZ-RhoGEF or other molecules mediate Rho activation with heightened intracellular Ca^2+^ levels in fibroblasts.

Besides the effectors we identified here, focal adhesion proteins, including focal adhesion kinase (FAK) and proteins of the FAK–Src signaling complex, are also known as crucial modulators participating in interactions with the extracellular matrix and the cytoskeleton(Parsons et al., 2010; Giannone et al., 2007; Gardel et al., 2010; Mitra et al., 2005). Intracellular forces generated by focal adhesion proteins promote rear retraction and the forward movement of the cell. The dynamic turnover of focal adhesions is spatiotemporally controlled by intracellular Ca^2+^ signaling(MacHacek et al., 2009; Mitra et al., 2005). In addition, calpains allow the degradation of FAPs, including FAK and Talin, in a Ca^2+^-dependent manner(Kerstein et al., 2017; Goll et al., 2003). Based on the extensive peripheral focal adhesions we observed in *Mfn2*-null MEFs, calpains or other focal adhesion proteins may also contribute to the “PAB” structure.

Neonates are susceptible to MFN2 defects (Filadi et al., 2018), and over 100 dominant mutations in the MFN2 gene have been reported in Charcot–Marie-Tooth disease type 2A (CMT2A) patients. However, how these mutations lead to disease is largely unknown, and there is currently no cure for this disease (Verhoeven et al., 2006; Calvo et al., 2009). MFN2 mutations are also associated with many other diseases, such as Alzheimer’s disease, Parkinson’s disease, obesity, and diabetes (Kim et al., 2017; Wang et al., 2009; Lee et al., 2012; Bach et al., 2003). One of the challenges in MFN2 research is that MFN2 plays multiple functional roles in cell signalings, such as regulating mitochondrial dynamics, transport, mtDNA stability, lipid metabolism, and cell survival. Both gain-of-function and loss-of-function mutations are reported in CMT2A patients. Some MFN2 mutations lead to fusion□incompetent mitochondria. However, some are fusion competent mutations (Strickland et al., 2014; Cartoni et al., 2010; Fissi et al., 2018; Franco et al., 2016; Rocha et al., 2018). It is also worth noting that MFN2 dysfunction preferentially impacts peripheral nerves instead of central nerves. These phenotypes are generally attributed to deficient mitochondrial trafficking and localization to the dendrites(Pareyson et al., 2015; Baloh et al., 2007). Our observations may provide a potential novel explanation of how MFN2 deficiency affects cell physiology by modulating the mechanosensitivity and cytoskeletal organization of the cells. It is possible that the MFN2 disease mutations also disrupt the mitochondria-ER tether and result in defects in cell cytoskeleton architecture, cell spreading, and migration, which cause the progression of the diseases. It is also possible that the same MFN2 mutation induces distinct signaling alterations in different cell types or diseases.

In summary, we characterized the alteration of the cytoskeleton and biophysical properties in MFN2-deficient cells and identified the detailed molecular mechanism. Our work provides insights into how MFN2 deficiency affects cell morphology and motility, specifically via the actin cytoskeleton.

## Materials and methods

### Cell culture

HEK293T (CRL-11268), wild-type (CRL-2991), *Mfn2*-null (CRL-2993), and *Mfn1*-null (CRL-2992) MEFs were from the American Type Culture Collection (ATCC, Manassas, VA, USA). GP2-293 cells were purchased from Takara Bio USA (#631458). All cells were maintained at 37°C with 5% CO_2_ in a Forma™ Steri-Cycle™ i160 CO_2_ Incubator (NC1207547, ThermoFisher Scientific). Cells were cultured in 10% FBS in DMEM with sodium bicarbonate. To obtain MFN2 reexpression cell line, introduce the ER-mito tethering structure, or express CaMKII-WT, CaMKII-DN, MLCK-CA, or MRLC-GFP, the MSCV-puro vector (Takara Bio USA, #634401) were used. We cloned the gene of interest into the MSCV vector and cotransfected the vector plasmid along with the envelope plasmid pVSVG, at a ratio of 1:1, into GP2-293 cells using Lipofectamin 3000 (Invitrogen L3000015). Virus supernatant was collected at both 48 hpt and 72 hpt and further concentrated with Lenti-X concentrator (Clotech 631232). MEF cells were transduced with concentrated retrovirus in a complete medium and then selected with 2 μg/ml puromycin (Gibco A1113803) starting the next day. The stable lines were generated after puromycin selection for one week. To generate ROCK or MLCK knocking down lines in *Mfn2*-null MEF cells, pLKO.1 lentiviral constructs with shRNAs were obtained from Sigma-Aldrich (shROCK: TRCN0000022903, shMLCK: TRCN0000024037), and SHC 003 was used as a nontargeting control. The plasmid of lentiviral constructs together with pCMV-dR8.2 dvpr (addgene #8455) and pCMV-VSV-G (addgene #8454), at a ratio of 10:7.5:2.5, were cotransfected into HEK293T cells with Lipofectamin 3000 (Invitrogen L3000015) to produce lentivirus. Virus supernatant was collected at both 48 hpt and 72 hpt, and further concentrated with Lenti-X concentrator (Clotech 631232) before transduction. 2 μg/ml puromycin (Gibco A1113803) was added into the complete medium on the next day for selection.

### Plasmids

In-Fusion cloning (In-Fusion HD Cloning Plus Kit, Clontech) was used to fuse the fragments with the linearized backbone. MSCV-puro was digested by Xho I and EcoR I. pLIX_403 plasmid was digested by NsiI and BamHI. The plasmids of Mfn1-Myc and Mfn2-Myc were gifts from David Chan (Addgene plasmid # 23212, #23213). The mito-GFP-ER plasmid was from the plasmid used in our lab (Addgene plasmid #160509). pLIX_403 was a gift from David Root (Addgene plasmid # 41395). GFP-C1-CAMKIIalpha and GFP-C1-CAMKIIalpha-K42R were gifts from Tobias Meyer (Addgene #21226, #21221). pTK91_GFP-MRLC2 was also from Addgene (Addgene #46355). The lentiviral backbone pLIX_403 was a gift from David Root (Addgene plasmid # 41395). pSLIK CA MLCK was a gift from Sanjay Kumar (Addgene plasmid # 84647). pYFP-Paxillin was a gift from Kenneth Yamada (Addgene #50543).

The In-Fusion primers are listed below: MSCV-mfn2 insert F: CACGATAATACCATGGGCCACCATGTCCCTGCTC MSCV-mfn2 insert R: TCTAGAGTCGCGGCCGCTTACTTGTACAGCTCGTCCATGCC MSCV-mfn1 insert R: TCGACTCTAGAGTCGCGGCCGCTTACTTGTACAGCTCGTCCATGCC Mfn2 into plix-Nsil-F: AAAACCCCGGTCCTATGCATATGTCCCTGCTCTTCTCTCGA Mfn2 into plix-BamHI-R: CCCCAACCCCGGATCCTTATCTGCTGGGCTGCAGGT Camk2a-MSCV-F: AATTAGATCTCTCGAGGCCACCATGGTGAGCAAGG Camk2a-MSCV-R: CTACCCGGTAGAATTCATTCGGCGAAGCAAGAGCG ER-mito F: AATTAGATCTCTCGAGATGGCAATCCAGTTGCGTTCG ER-mito R: ATTTACGTAGCGGCCGCTTAAGATACATTGATGAGTTTGG MRLC-GFP F: AATTAGATCTCTCGAGGCCACCATGGTGAGCAAGG MRLC-GFP R: CTACCCGGTAGAATTCGCCCGCGGTCAGTCATCTTTG MLCK-CA F: ATTAGATCTCTCGAGACTAGTCGACTGGATCC MLCK-CA R: CCGGTAGAATTCAGATCTTGGGTGGGTTAATTAA

### Chemicals

MEF cells were treated with 1% DMSO, 20 μM BAPTA (Cayman Chemical), Y29632 5μM (Cayman Chemical), 50 μM CK666 (Cayman Chemical), STO-609 acetate (Biotechne, #1551), A23187 (Cayman Chemical), 4 μM Blebbistatin (Cayman Chemical), 40 uM AIP (R&D SYSTEMS #5959/1), 50 μM CAS 1090893 (Millipore, #553511), 0.1 μg/ml RhoA inhibitor-I (Cytoskeleton, Inc., #CT-04), 2 μM FK-506 (Cayman No. 10007965), 300 nM FAK14 (Cayman, #14485), 10 μM BMS-5 (Cayman Chemical, #21072) or 2 μM ML-7 (Cayman Chemical, #11801) overnight during time-lapse imaging for cell random migration or followed by immunostaining.

### Western Blot

Total protein was isolated from cells using RIPA buffer containing 25 mM Tris-HCl (pH 8.0), 150 mM NaCl, 1 mM EDTA, 0.5% NP-40, 0.5% sodium deoxycholate and, 0.1% sodium dodecyl sulfate (SDS). For samples containing phosphor proteins to probe, 20mM sodium fluoride (NaF), 1mM sodium orthovanadate (Na2VO3), 10 mM beta glycerophosphate were added to the RIPA lysis buffer. Protein concentrations were determined using the Precision Red Advanced Protein Assay Reagent (Cytoskeleton ADV02). Extracted proteins (25-35 μg) were separated by 8-12% SDS-PAGE and transferred onto polyvinylidene difluoride membranes (PVDF, BioRad). Membranes were blocked for ~30 min in PBST (PBS and 0.1% Tween 20) with 5% fat-free milk. After blocking, membranes were incubated with primary antibodies diluted 1:1,000 in 1% BSA at 4°C overnight, and secondary antibodies diluted 1:10,000 in PBST at room temperature for 1 h. Odyssey (LI-Cor) was used to image membranes. Immunodetection of the pulldown samples was performed using enhanced Western Blotting Chemiluminescence Luminol Reagent (Santa Cruz Biotechnology, cat#sc-2048) and detected with a FluorChem R System (Proteinsimple). Image Studio 5.0 was used to quantify and analyze the results. Primary antibodies anti-Mfn2 (Cell Signaling 9482S), anti-Mfn1 (Abcam, ab126575), anti-pan-CaMK(Cell Signaling #3362), anti-phosphor-CaMKII(Thr286) (Cell Signaling #12716), anti-phospho Myosin Light Chain 2 (Cell Signaling #3671), anti-Myosin Light Chain 2(Cell Signaling #3672), anti-phospho-PAK (Cell signaling #2605S), anti-PAK (Cell signaling #2604) and secondary antibody HRP AffiniPure goat anti-rabbit IgG (Jackson ImmunoReseach, #111-035-003), goat anti-mouse IgG Alexa Fluor 680 (Invitrogen, #A28183), goat anti-rabbit IgG Alexa Fluor Plus 800 (Invitrogen, #A32735).

### Rac-GTP and RhoA-GTP pulldown assay

PAK-GST-coated beads (Cytoskeleton BK035) and Rhotekin-RBD-coated beads(Cytoskeleton BK036) were used to isolate active Rac and RhoA from the whole-cell lysate. MEF cells were serum starved with DMEM medium lacking FBS overnight in the incubator at 70%-80% confluency. After starvation, PDGF-BB was then added to the cells at a final concentration of 25μM. Then cells were lysed with ice-cold lysis buffer at indicated time points and collected by scrapples. 15μg PAK-GST beads or 50μg Rhotekin-RBD-coated beads were mixed with each sample and incubated at 4°C for 1 h. Protein beads were washed and processed for western blot.

### Immunostaining and confocal imaging

For immunofluorescent staining, MEF cells with or without drug treated overnight were plated onto coverslips and incubated overnight at 37°C, then fixed with 4% Paraformaldehyde (PFA) solution in PBS for 15 mins at room temperature. Cells were permeabilized in PBS with 0.1% Triton X-100 and 3% fatty-acid-free BSA for 1 hour, then incubated with phalloidin -AlexaFluor 488 (Invitrogen A12379) or primary antibodies diluted 1:100 in 3% BSA overnight at 4°C. After washing with PBS three times, the cells were then stained with secondary antibodies diluted 1:500 in 3% BSA and DAPI (Invitrogen D3571) for 1 h at room temperature. After washing with PBS three times, the coverslips were mounted on glass slides with the mounting medium (Vector Laboratories H-1000). Images were acquired using an N-STORM/N-SIM TIRF microscope (Nikon) with a 1.49/60x Apo TIRF oil objective and a laser-scanning confocal microscope (LSM 800, Zeiss) with a 1.4/63x oil immersion objective lens. Images were processed and analyzed with ImageJ. Primary antibodies anti-Mfn2 (Cell Signaling 9482S), anti-Paxillin (Invitrogen AHO0492), anti-phospho Myosin Light Chain 2 (Cell Signaling 3671) and secondary antibody anti-rabbit AlexaFluor 488 (Invitrogen A-21441), anti-mouse AlexaFluor 568 (Invitrogen A-11004) were used. The cell area and circularity were measured using ImageJ and plotted in Prism 6.0 (GraphPad).

### 2D migration live imaging

MEF cells were first trypsinized and replated onto fibrinogen-coated μ-slide 8 well plates (ibidi 80826) at a density of ~10000 cells per well with a complete medium. Time-lapse images were acquired using BioTek Lionheart FX Automated Microscope with 20x phase lens at 10 min intervals of ~18 h at 37°C with 5% CO_2_. The velocity of MEFs was measured using ImageJ with the MTrackJ plugin and plotted in Prism 6.0 (GraphPad).

### Ca^2+^ measurement

Fluo-4 Calcium Imaging Kit (Invitrogen F10489) was used for cytosolic Ca^2+^ measurement in MEFs. MEF cells were incubated with PowerLoad solution and Fluo-4 dye at 37 °C for 15 min and then at room temperature for 15 min. After incubation, cells were washed with PBS for one time. Time-lapse green fluorescence images were obtained with AXIO Zoom V16 microscope (Zeiss) at 1 min interval of 25 mins. 50μl of 2mM PDGF-BB(sigma, #P4056) was added to cells right after the first image was taken. The fold change of the fluorescence intensity was normalized to that of the first image. The fluorescence intensity was measured using ImageJ and plotted in Prism 6.0 (GraphPad).

### Single-molecule super-resolution imaging

Single-molecule super-resolution imaging was performed on a custom-built setup on an Olympus IX-73 microscope stand (Olympus America, IX-73) equipped with a 100×/1.35-NA silicone-oil-immersion objective lens (Olympus America, UPLSAPO100XS) and a PIFOC objective positioner (Physik Instrumente, ND72Z2LAQ). Samples were excited by a 642 nm laser (MPB Communications, 2RU-VFL-P-2000-642-B1R), which passed through an acoustic-optic tunable filter (AA Opto-electronic, AOTFnC-400.650-TN) for power modulation. The excitation light was focused on the pupil plane of the objective lens after passing through a filter cube holding a quadband dichroic mirror (Chroma, ZT405/488/561/647rpc). The fluorescent signal was magnified by relay lenses arranged in a 4f alignment to a final magnification of ~54 and then split with a 50/50 non-polarising beam splitter (Thorlabs, BS016). The split fluorescent signals were delivered by two mirrors onto a 90° specialty mirror (Edmund Optics, 47-005), axially separated by 590 nm in the sample plane, and then projected on an sCMOS camera (Hamamatsu, Orca-Flash4.0v3) with an effective pixel size of 119 nm. A bandpass filter (Semrock, FF01-731/137-25) was placed before detection. The imaging system was controlled by custom-written LabVIEW (National Instruments) programs.

Before imaging, the coverslip with cells on top was placed on a custom-made holder. 100 μL of imaging buffer (10% (w/v) glucose in 50 mM Tris, 50 mM NaCl, 10 mM ß-Mercaptoethylamine hydrochloride (M6500, Sigma-Aldrich), 50 mM 2-Mercaptoethanol (M3148, Sigma-Aldrich), 2 mM cyclooctatetraene (138924, Sigma-Aldrich), 2.5 mM protocatechuic acid (37580, Sigma-Aldrich) and 50 nM protocatechuate 3,4-Dioxygenase (P8279, Sigma-Aldrich), pH 8.0) were added on top of the coverslip. Then another coverslip was placed on top of the imaging buffer. This coverslip sandwich was then sealed with two-component silicon dental glue (Dental-Produktions und Vertriebs GmbH, picodent twinsil speed 22).

Following the previous procedure (Xu et al., 2020), the sample was first excited with the 642-nm laser at a low intensity of ~50 W/cm^2^ to find a region of interest. Before fluorescence imaging, bright-field images of this region were recorded over an axial range from −1 to +1 μm with a step size of 100 nm as reference images for focus stabilization. Single-molecule blinking data were then collected at a laser intensity of 2–6 kW/cm^2^ and a frame rate of 50 Hz. Imaging was conducted for ~30 cycles with 2,000 frames per cycle. Single-molecule localization was performed as described previously (Xu et al., 2020).

### Seahorse mitochondrial respiration analysis

Mitochondrial respiration was measured with Seahorse XFe24 Analyzer (Agilent Technologies) according to the manual of Seahorse XF Cell Mito Stress Test Kit (Agilent Technologies, cat#103015-100). Briefly, MEF cells were plated on the XF24 cell culture microplate at a density of 50,000 cells per well. The seahorse sensor cartridge was hydrated with calibrant in a non-CO_2_ incubator at 37°C overnight 1 day before measurement. On the day of measurements, cells were washed once and incubated in Seahorse XF base medium (pH7.4, Agilent Technologies, cat#103334-100) supplemented with 1 mM sodium pyruvate, 2 mM glutamine, and 5.5 mM glucose. Cells were equilibrated at 37°C in a non-CO_2_ incubator for 1 hour. The oxygen consumption rate was monitored at the basal state and after sequential injection of the mitochondrial compounds oligomycin (1 μg/mL), FCCP (1 μM), and Rotenone/antimycin A (both 1 μM) to induce mitochondrial stress. All mitochondrial respiration rates were generated and automatically calculated by the Seahorse Wave software with normalization to the cellular protein contents. Cellular protein contents were determined by the sulforhodamine B (SRB) assay as described (Vichai and Kirtikara, 2006).

### Traction Force Microscopy and analysis

Traction force microscopy was performed as described previously (Huang et al., 2019; Sala and Oakes, 2021; Aratyn-Schaus et al., 2010). Briefly, 22 x 30mm #1.5 glass coverslips were activated by incubating with a 2% solution of 3-aminopropyltrimethyoxysilane (313255000, Acros Organics) diluted in isopropanol, followed by fixation in 1% glutaraldehyde (16360, Electron Microscopy Sciences) in ddH_2_0. Polyacrylamide gels (shear modulus: 16 kPa—final concentrations of 12% acrylamide [1610140, Bio-Rad] and 0.15% bis-acrylamide [1610142, Bio-Rad]) were embedded with 0.04-μm fluorescent microspheres (F8789, Invitrogen) and ~6 mg/mL acryloyl-X, SE (6-((acryloyl)amino)hexanoic acid)-labelled fibronectin (A20770, ThermoFisher Scientific; FC010, EMD Millipore), and polymerized on activated glass coverslips for 1 hr at room temperature. After polymerization, gels were rehydrated for 1 hr in deionized H_2_O, before seeding 1.0× 10^5^ cells on each gel in a 60 mm cell culture-treated petri dish. Cells were allowed to spread overnight, and the next day SPY555-actin dye (SC202, SPIROCHROME) was added ~60 min before imaging. Images were taken of both the cells and underlying fluorescent beads. Following imaging, cells were removed from the gel by adding 0.025% SDS, and a reference image of the fluorescent beads in the unstrained gel was taken.

Analysis of traction forces was performed using code written in Python according to previously described approaches (Sabass et al., 2008). Prior to processing, the reference bead image was aligned to the bead image with the cell attached. Displacements in the beads were calculated using an optical flow algorithm in OpenCV (Open Source Computer Vision Library, https://github/itseez/opencv) with a window size of 16 pixels. Traction stresses were calculated using the FTTC approach(Butler et al., 2002; Huang et al., 2019) as previously described, with a regularization parameter of 9.34× 10^-9^. The strain energy was calculated by summing one-half the product of the strain and traction vectors in the region under the cell (Oakes et al., 2014b) and normalized by the cell area as measured using the SPY555-actin image of the cell. Cells with the residual energy of ≥ 20% were excluded from the data set. N= 4 with ≥ 9 cells per biological repeat.

### Substrate stiffness assay

Polyacrylamide gels with uniform stiffness were prepared as previously described(Efremov et al., 2022). Briefly, 50 mm glass bottom dishes (WPI, USA) were activated with 0.1 M NaOH, 4% (v/v) APTES ((3-Aminopropyl)triethoxysilane, Sigma-Aldrich, USA) and 1% (v/v) glutaraldehyde (Sigma-Aldrich, USA) in 1X PBS. To prepare a gel with a specific stiffness, different concentrations of acrylamide (40%, Sigma-Aldrich, USA) and bis-acrylamide (1%, Sigma-Aldrich, USA) were mixed with PBS 1X and 0.5 % (w/v) Irgacure 2959 (0.5% w/v, (2-Hydroxy-4’-(2-hydroxyethoxy)-2-methylpropiophenone, Sigma-Aldrich, USA). Later, the gel solution was incubated at 37°C overnight and degassed for 30 minutes at room temperature. Next, to prepare one gel, 120 ul of the gel solution were poured into the center of an activated glass bottom dish and covered with 22*22 mm coverglass previously chloro-silanated with DCDMS (Dichlorodimethylsilane, Sigma-Aldrich, USA). Then, the dish was placed in a UV transilluminator for 10 minutes. Final concentrations of acrylamide (4%) and bis-acrylamide (0.2%, w/v) were chosen to prepare PAA gels with 1.67, 4.47, 10.61, 16.7 and 19.66kPa Young’s modulus. The ratio is shown below:

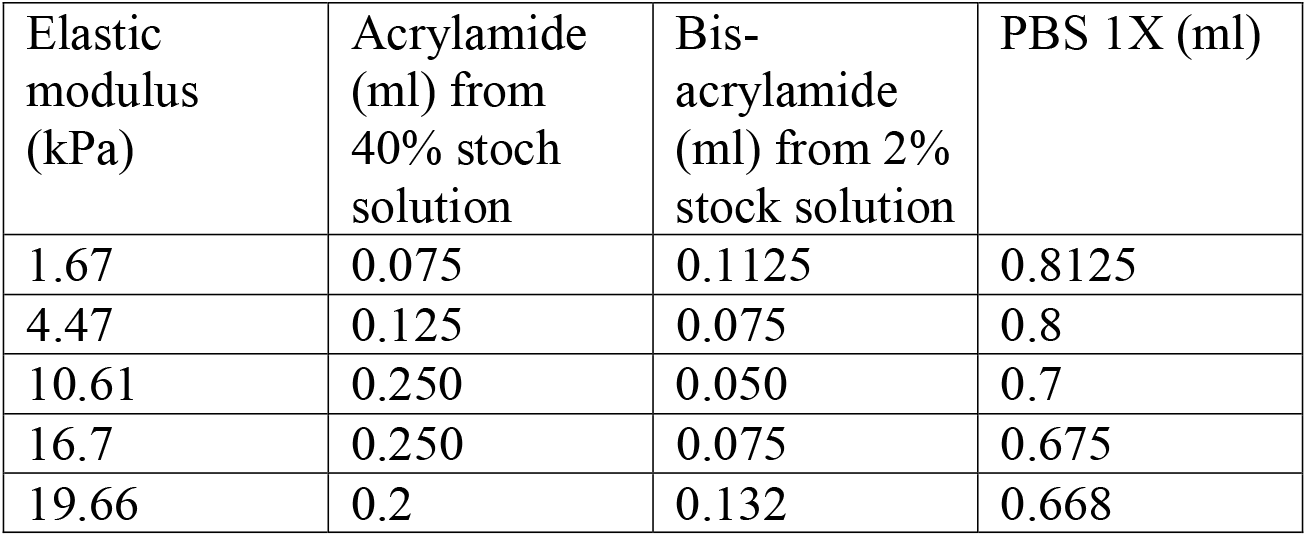

### Atomic force microscopy

An Asylum Research MFP3d Bio AFM system (Santa Barbara, CA) was used to measure the effective modulus of *wt* and *Mfn2*-null MEFs, seeded in 60 mm polystyrene Petri dishes. A Nanoandmore CP-qp-SCONT-SiO-B-5 colloidal probe cantilever (Watsonville, CA) with 3.5-μm probe diameter and 0.01 N/m nominal stiffness was used for these experiments. The optical lever sensitivity of the cantilever was calibrated using a static force-displacement curve, performed on the part of the polystyrene petri dish not covered by the cells. The cantilever stiffness was determined to be 0.00664 N/m using the thermal tuning method in the air (Hutter and Bechhoefer, 1998). AFM indentation experiments were performed using cells in phosphate-buffered saline (PBS) solution in a 6-cm petri dish maintained at a constant temperature of 37°C using a petri dish heater. For imaging, a sufficient amount of PBS was used to maintain cell viability, and a 100μL drop of PBS was placed on the AFM cantilever tip to avoid the formation of air bubbles between the cantilever holder and the sample. Force-displacement curves were acquired for each cell by indenting them at the central region of the cell, close to the nucleus. The force spectroscopy experiments were performed for a single force cycle by setting trigger points on the cantilever deflection (u) and tip velocity. A time gap of > 45 sec was incorporated between the indentation experiments within the same cell to account for stress relaxation. The force-displacement curve corresponding to the approach of the cantilever tip towards the substrate for each location was considered to quantify the effective cell modulus using the material properties of the cantilever, and Hertzian contact mechanics model for a spherical indenting a flat plane (Hertz 1881, Guo and Akhremitchev, 2006) According to this model, the force measured during cell indentation (F) is related to the cantilever indentation, δ, as (Johnson, 1985)

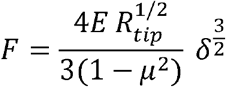

Where μ is the Poisson’s ratio, E is the effective cell modulus, and R_tip_ is the radius of the spherical cantilever tip. The cell is assumed to be incompressible with μ = 0.5. The force curves were delineated into a region prior to the contact point of the cantilever tip with the cell and the region after contact. Force data post contact with the cell was used to calculate the effective cell modulus.

### FLIM imaging

Frequency-domain fluorescence lifetime imaging microscopy (FLIM) measurements were performed using a Nikon TE2000 confocal microscope with a 60×/1.2NA water immersion objective equipped with an Alba FastFLIM system (Sun et al., 2011). Specifically, cells stained with Flipper-TR (Cytoskeleton, CY-SC020) were excited using a 488nm pulsed laser with a modulation frequency of 20MHz and imaged through a 595/40nm bandpass filter followed by MPD APD detectors. After image collection, bi-exponential fitting of FLIM images was performed using VistaVision software (ISS) to obtain fluorescent lifetimes (*τ*_1_ and *τ*_2_) of each pixel. Only the longest lifetime component *τ*_1_) was used to represent relative membrane tension as described previously (Colom et al., 2018).

### Statistical analysis

Statistical analysis was performed with Prism 6 (GraphPad). An unpaired two-tailed Student’s *t-* test or one-way ANOVA was used to determine the statistical significance of differences between groups. A *P*-value less than 0.05 was considered statistically significant.

Individual *P* values are indicated in the figures, with no data points excluded from statistical analysis. One representative experiment of at least 3 independent repeats is shown.

## Acknowledgments

The work was supported by the National Institutes of Health [R35GM119787 to DQ], and [P30CA023168 to Purdue Center for Cancer Research] for shared resources. This work is based upon efforts supported by EMBRIO Institute, contract #2120200, a National Science Foundation (NSF) Biology Integration Institute (to DC). YW is supported by Bisland Fellowship, Purdue University.

## Author contributions

YW and DQ designed the research and wrote the manuscript. YW, LT, FX, AC, HZ, LC, RW. JC performed experiments and analyzed the data. SK, DS, CY, DC. FH, PO and DQ designed experiments and supervised the research. All authors read and approved the manuscript.

## Competing interests

The authors declare no competing interests.

## Data availability statement

Plasmids will be available on Addgene.

## Notes

### Competing Interest Statement

The authors have declared no competing interest.

### Summary of Updates

Errors in supplemental files removed.

## References

Aguilar-Cuenca, R., A. Juanes-García, and M. Vicente-Manzanares. 2014. Myosin II in mechanotransduction: master and commander of cell migration, morphogenesis, and cancer. Cell Mol Life Sci. 71:479–492. doi:10.1007/S00018-013-1439-5.

Amano, M., M. Ito, K. Kimura, Y. Fukata, K. Chihara, T. Nakano, Y. Matsuura, and K. Kaibuchi. 1996. Phosphorylation and activation of myosin by Rho-associated kinase (Rhokinase). J Biol Chem. 271:20246–20249. doi:10.1074/JBC.271.34.20246.

Aratyn-Schaus, Y., P.W. Oakes, J. Stricker, S.P. Winter, and M.L. Gardel. 2010. Preparation of Complaint Matrices for Quantifying Cellular Contraction. J Vis Exp. 46. doi:10.3791/2173.

Bach, D., S. Pich, F.X. Soriano, N. Vega, B. Baumgartner, J. Oriola, J.R. Daugaard, J. Lloberas, M. Camps, J.R. Zierath, R. Rabasa-Lhoret, H. Wallberg-Henriksson, M. Laville, M. Palacín, H. Vidal, F. Rivera, M. Brand, and A. Zorzano. 2003. Mitofusin-2 determines mitochondrial network architecture and mitochondrial metabolism. A novel regulatory mechanism altered in obesity. J Biol Chem. 278:17190–17197. doi:10.1074/JBC.M212754200.

Baloh, R.H., R.E. Schmidt, A. Pestronk, and J. Milbrandt. 2007. Altered axonal mitochondrial transport in the pathogenesis of Charcot-Marie-Tooth disease from mitofusin 2 mutations. J Neurosci. 27:422–430. doi:10.1523/JNEUROSCI.4798-06.2007.

Báthori, G., G. Csordás, C. Garcia-Perez, E. Davies, and G. Hajnóczky. 2006. Ca2+-dependent control of the permeability properties of the mitochondrial outer membrane and voltage-dependent anion-selective channel (VDAC). Journal of Biological Chemistry. 281:17347–17358. doi:10.1074/jbc.M600906200.

Bennett, J., and A. Weeds. 1986. CALCIUM AND THE CYTOSKELETON. Br Med Bull.42:385–390. doi:10.1093/OXFORDJOURNALS.BMB.A072156.

de Brito, O.M., and L. Scorrano. 2008. Mitofusin 2 tethers endoplasmic reticulum to mitochondria. Nature 2008 456:7222. 456:605–610. doi:10.1038/nature07534.

de Bruyn, P.P.H. 1946. The amoeboid movement of the mammalian leukocyte in tissue culture. Anat Rec. 95:177–191. doi:10.1002/AR.1090950209.

Burridge, K., and M. Chrzanowska-Wodnicka. 2003. FOCAL ADHESIONS, CONTRACTILITY, AND SIGNALING. http://dx.doi.org/10.1146/annurev.cellbio.12.1.463. 12:463–519. doi:10.1146/ANNUREV.CELLBIO.12.1.463.

Butler, J.P., I.M. Toli-Nørrelykke, B. Fabry, and J.J. Fredberg. 2002. Traction fields, moments, and strain energy that cells exert on their surroundings. Am J Physiol Cell Physiol. 282. doi:10.1152/AJPCELL.00270.2001.

Calvo, J., B. Funalot, R.A. Ouvrier, L. Lazaro, A. Toutain, P. de Mas, P. Bouche, B. Gilbert-Dussardier, M.C. Arne-Bes, J.P. Carrière, H. Journel, M.C. Minot-Myhie, C. Guillou, K. Ghorab, L. Magy, F. Sturtz, J.M. Vallat, and C. Magdelaine. 2009. Genotype-Phenotype Correlations in Charcot-Marie-Tooth Disease Type 2 Caused by Mitofusin 2 Mutations. Arch Neurol. 66:1511–1516. doi:10.1001/ARCHNEUROL.2009.284.

Campello, S., R.A. Lacalle, M. Bettella, S. Mañes, L. Scorrano, and A. Viola. 2006. Orchestration of lymphocyte chemotaxis by mitochondrial dynamics. Journal of Experimental Medicine. 203:2879–2886. doi:10.1084/JEM.20061877/VIDEO-7.

Cartoni, R., E. Arnaud, J.J. Médard, O. Poirot, D.S. Courvoisier, R. Chrast, and J.C. Martinou. 2010. Expression of mitofusin 2(R94Q) in a transgenic mouse leads to Charcot-Marie-Tooth neuropathy type 2A. Brain. 133:1460–1469. doi:10.1093/BRAIN/AWQ082.

Chen, H., S.A. Detmer, A.J. Ewald, E.E. Griffin, S.E. Fraser, and D.C. Chan. 2003a. Mitofusins Mfn1 and Mfn2 coordinately regulate mitochondrial fusion and are essential for embryonic development. Journal of Cell Biology. 160:189–200. doi:10.1083/jcb.200211046.

Cipolat, S., O.M. de Brito, B. Dal Zilio, and L. Scorrano. 2004. OPA1 requires mitofusin 1 to promote mitochondrial fusion. Proc Natl Acad Sci USA. 101:15927–15932. doi:10.1073/PNAS.0407043101/SUPPL_FILE/07043FIG6.JPG.

Clapham, D.E. 2007a. Calcium signaling. Cell. 131:1047–1058. doi:10.1016/J.CELL.2007.11.028.

Colom, A., E. Derivery, S. Soleimanpour, C. Tomba, M.D. Molin, N. Sakai, M. González-Gaitán, S. Matile, and A. Roux. 2018. A fluorescent membrane tension probe. Nature Chemistry 2018 10:11. 10:1118–1125. doi:10.1038/s41557-018-0127-3.

Denisenko, T. v., A.S. Gorbunova, and B. Zhivotovsky. 2019. Mitochondrial Involvement in Migration, Invasion and Metastasis. Front Cell Dev Biol. 7:355. doi:10.3389/FCELL.2019.00355/XML/NLM.

Derewenda, U., A. Oleksy, A.S. Stevenson, J. Korczynska, Z. Dauter, A.P. Somlyo, J. Otlewski, A. v. Somlyo, and Z.S. Derewenda. 2004. The crystal structure of RhoA in complex with the DH/PH fragment of PDZRhoGEF, an activator of the Ca(2+) sensitization pathway in smooth muscle. Structure. 12:1955–1965. doi:10.1016/J.STR.2004.09.003.

Discher, D.E., P. Janmey, and Y.L. Wang. 2005. Tissue cells feel and respond to the stiffness of their substrate. Science (1979). 310:1139–1143. doi:10.1126/SCIENCE.1116995/ASSET/9FD0A405-058F-4F6E-B5B4-F55BB0CF5F4A/ASSETS/GRAPHIC/310_1139_F4.JPEG.

Dorn, G.W. 2020. Mitofusins as mitochondrial anchors and tethers. J Mol Cell Cardiol. 142:146–153. doi:10.1016/j.yjmcc.2020.04.016.

Efremov, Y.M., D.M. Suter, P.S. Timashev, and A. Raman. 2022. 3D nanomechanical mapping of subcellular and sub-nuclear structures of living cells by multi-harmonic AFM with long-tip microcantilevers. Scientific Reports 2022 12:1. 12:1–11. doi:10.1038/s41598-021-04443-w.

Efremov, Y.M., M. Velay-Lizancos, C.J. Weaver, A.I. Athamneh, P.D. Zavattieri, D.M. Suter, and A. Raman. 2019. Anisotropy vs isotropy in living cell indentation with AFM. Scientific Reports 2019 9:1. 9:1–12. doi:10.1038/s41598-019-42077-1.

Filadi, R., E. Greotti, G. Turacchio, A. Luini, T. Pozzan, and P. Pizzo. 2015a. Mitofusin 2 ablation increases endoplasmic reticulum-mitochondria coupling. Proc Natl Acad Sci U S A.112:E2174–E2181. doi:10.1073/pnas.1504880112.

Filadi, R., Di. Pendin, and P. Pizzo. 2018. Mitofusin 2: from functions to disease. Cell Death & Disease 2018 9:3. 9:1–13. doi:10.1038/s41419-017-0023-6.

Fissi, N. el, M. Rojo, A. Aouane, E. Karatas, G. Poliacikova, C. David, J. Royet, and T. Rival. 2018. Mitofusin gain and loss of function drive pathogenesis in Drosophila models of CMT2A neuropathy. EMBO Rep. 19. doi:10.15252/EMBR.201745241.

Fleming, I.N., C.M. Elliott, F.G. Buchanan, C.P. Downes, and J.H. Exton. 1999. Ca2+/calmodulin-dependent protein kinase II regulates Tiam1 by reversible protein phosphorylation. J Biol Chem. 274:12753–12758. doi:10.1074/JBC.274.18.12753.

Franco, A., R.N. Kitsis, J. A. Fleischer, E. Gavathiotis, O.S. Kornfeld, G. Gong, N. Biris, A. Benz, N. Qvit, S.K. Donnelly, Y. Chen, S. Mennerick, L. Hodgson, D. Mochly-Rosen, and G.W. Dorn. 2016. Correcting mitochondrial fusion by manipulating mitofusin conformations. Nature. 540:74–79. doi:10.1038/NATURE20156.

Gardel, M.L., I.C. Schneider, Y. Aratyn-Schaus, and C.M. Waterman. 2010. Mechanical Integration of Actin and Adhesion Dynamics in Cell Migration. Annu Rev Cell Dev Biol. 26:315. doi:10.1146/ANNUREV.CELLBIO.011209.122036.

Giannone, G., B.J. Dubin-Thaler, O. Rossier, Y. Cai, O. Chaga, G. Jiang, W. Beaver, H.G. Döbereiner, Y. Freund, G. Borisy, and M.P. Sheetz. 2007. Lamellipodial actin mechanically links myosin activity with adhesion site formation. Cell. 128:561. doi:10.1016/J.CELL.2006.12.039.

Goll, D.E., V.F. Thompson, H. Li, W. Wei, and J. Cong. 2003. The calpain system. Physiol Rev.83:731–801. doi:10.1152/PHYSREV.00029.2002.

Guo, S., and B.B. Akhremitchev. 2006. Packing density and structural heterogeneity of insulin amyloid fibrils measured by AFM nanoindentation. Biomacromolecules. 7:1630–1636. doi:10.1021/BM0600724/SUPPL_FILE/BM0600724SI20060313_041959.PDF.

Hall, A. 1998a. Rho GTPases and the Actin Cytoskeleton. Science (1979). 279:509–514. doi:10.1126/SCIENCE.279.5350.509.

Hall, A. 2005. Rho GTPases and the control of cell behaviour. Biochem Soc Trans.33:891–895. doi:10.1042/BST20050891.

Huang, Y., C. Schell, T.B. Huber, A.N. Şimşek, N. Hersch, R. Merkel, G. Gompper, and B. Sabass. 2019. Traction force microscopy with optimized regularization and automated Bayesian parameter selection for comparing cells. Sci Rep. 9. doi:10.1038/S41598-018-36896-X.

Hudmon, A., and H. Schulman. 2002. Structure-function of the multifunctional Ca2+/calmodulin-dependent protein kinase II. Biochem J. 364:593–611. doi:10.1042/BJ20020228.

Hutter, J.L., and J. Bechhoefer. 1998. Calibration of atomic force microscope tips. Review of Scientific Instruments. 64:1868. doi:10.1063/1.1143970.

Ikebe, M., and D.J. Hartshorne. 1985. Proteolysis of Smooth Muscle Myosin by Staphylococcus aureus Protease: Preparation of Heavy Meromyosin and Subfragment 1 with Intact 20000-Dalton Light Chains. Biochemistry. 24:2380–2387. doi:10.1021/bi00330a038.

Johnson, K.L. 1985. Contact Mechanics. doi:10.1017/CBO9781139171731.

Kaibuchi, K., S. Kuroda, and M. Amano. 1999. Regulation of the cytoskeleton and cell adhesion by the Rho family GTPases in mammalian cells. Annu Rev Biochem.68:459–486. doi:10.1146/ANNUREV.BIOCHEM.68.1.459.

Kamm, K.E., and J.T. Stull. 1985. The function of myosin and myosin light chain kinase phosphorylation in smooth muscle. Annu Rev Pharmacol Toxicol. 25:593–620. doi:10.1146/ANNUREV.PA.25.040185.003113.

Kassianidou, E., J.H. Hughes, and S. Kumar. 2017. Activation of ROCK and MLCK tunes regional stress fiber formation and mechanics via preferential myosin light chain phosphorylation. Mol Biol Cell. 28:3832–3843. doi:10.1091/MBC.E17-06-0401/ASSET/IMAGES/LARGE/3832FIG8.TIF.GZ.JPEG.

Katoh, K., Y. Kano, M. Amano, K. Kaibuchi, and K. Fujiwara. 2001a. Stress fiber organization regulated by MLCK and Rho-kinase in cultured human fibroblasts. Am J Physiol Cell Physiol. 280. doi:10.1152/AJPCELL.2001.280.6.C1669.

Katoh, K., Y. Kano, M. Amano, H. Onishi, K. Kaibuchi, and K. Fujiwara. 2001b. Rho-Kinase–Mediated Contraction of Isolated Stress Fibers. J Cell Biol. 153:569. doi:10.1083/JCB.153.3.569.

Kerstein, P.C., K.M. Patel, and T.M. Gomez. 2017. Calpain-Mediated Proteolysis of Talin and FAK Regulates Adhesion Dynamics Necessary for Axon Guidance. Journal of Neuroscience. 37:1568–1580. doi:10.1523/JNEUROSCI.2769-16.2016.

Kim, Y.J., J.K. Park, W.S. Kang, S.K. Kim, C. Han, H.R. Na, H.J. Park, J.W. Kim, Y.Y. Kim, M.H. Park, and J.W. Paik. 2017. Association between Mitofusin 2 Gene Polymorphisms and Late-Onset Alzheimer’s Disease in the Korean Population. Psychiatry Investig. 14:81–85. doi:10.4306/PI.2017.14.1.81.

Kornmann, B., E. Currie, S.R. Collins, M. Schuldiner, J. Nunnari, J.S. Weissman, and P. Walter. 2009. An ER-mitochondria tethering complex revealed by a synthetic biology screen. Science. 325:477–481. doi:10.1126/SCIENCE.1175088.

Lamb, M.C., C.P. Kaluarachchi, T.I. Lansakara, S.Q. Mellentine, Y. Lan, A. v. Tivanski, and T.L. Tootle. 2021. Fascin limits myosin activity within drosophila border cells to control substrate stiffness and promote migration. Elife. 10. doi:10.7554/ELIFE.69836.

Lämmermann, T., B.L. Bader, S.J. Monkley, T. Worbs, R. Wedlich-Söldner, K. Hirsch, M. Keller, R. Förster, D.R. Critchley, R. Fässler, and M. Sixt. 2008. Rapid leukocyte migration by integrin-independent flowing and squeezing. Nature. 453:51–55. doi:10.1038/NATURE06887.

Lauffenburger, D.A., and A.F. Horwitz. 1996. Cell migration: a physically integrated molecular process. Cell. 84:359–369. doi:10.1016/S0092-8674(00)81280-5.

Lee, S., F.H. Sterky, A. Mourier, M. Terzioglu, S. Cullheim, L. Olson, and N.G. Larsson. 2012. Mitofusin 2 is necessary for striatal axonal projections of midbrain dopamine neurons. Hum Mol Genet. 21:4827–4835. doi:10.1093/HMG/DDS352.

Lo, C.M., H.B. Wang, M. Dembo, and Y.L. Wang. 2000. Cell movement is guided by the rigidity of the substrate. Biophys J. 79:144–152. doi:10.1016/S0006-3495(00)76279-5.

MacHacek, M., L. Hodgson, C. Welch, H. Elliott, O. Pertz, P. Nalbant, A. Abell, G.L. Johnson, K.M. Hahn, and G. Danuser. 2009. Coordination of Rho GTPase activities during cell protrusion. Nature. 461:99–103. doi:10.1038/NATURE08242.

Maianski, N.A., F.P.J. Mul, J.D. van Buul, D. Roos, and T.W. Kuijpers. 2002. Granulocyte colony-stimulating factor inhibits the mitochondria-dependent activation of caspase-3 in neutrophils. Blood. 99:672–679. doi:10.1182/BLOOD.V99.2.672.

Mitra, S.K., D.A. Hanson, and D.D. Schlaepfer. 2005. Focal adhesion kinase: in command and control of cell motility. Nature Reviews Molecular Cell Biology 2005 6:1. 6:56–68. doi:10.1038/nrm1549.

Naon, D., M. Zaninello, M. Giacomello, T. Varanita, F. Grespi, S. Lakshminaranayan, A. Serafini, M. Semenzato, S. Herkenne, M.I. Hernández-Alvarez, A. Zorzano, D. de Stefani, G.W. Dorn, and L. Scorrano. 2016. Critical reappraisal confirms that Mitofusin 2 is an endoplasmic reticulum-mitochondria tether. Proc Natl Acad Sci U S A. 113:11249–11254. doi:10.1073/pnas.1606786113.

Nobes, C.D., and A. Hall. 1995. Rho, rac, and cdc42 GTPases regulate the assembly of multimolecular focal complexes associated with actin stress fibers, lamellipodia, and filopodia. Cell. 81:53–62. doi:10.1016/0092-8674(95)90370-4.

Nobes, C.D., and A. Hall. 1999. Rho GTPases Control Polarity, Protrusion, and Adhesion during Cell Movement. J Cell Biol. 144:1235. doi:10.1083/JCB.144.6.1235.

Oakes, P.W., S. Banerjee, M.C. Marchetti, and M.L. Gardel. 2014a. Geometry Regulates Traction Stresses in Adherent Cells. Biophys J. 107:825. doi:10.1016/J.BPJ.2014.06.045.

Okabe, T., T. Nakamura, Y.N. Nishimura, K. Kohu, S. Ohwada, Y. Morishita, and T. Akiyama. 2003. RICS, a novel GTPase-activating protein for Cdc42 and Rac1, is involved in the beta-catenin-N-cadherin and N-methyl-D-aspartate receptor signaling. J Biol Chem. 278:9920–9927. doi:10.1074/JBC.M208872200.

Pareyson, D., P. Saveri, A. Sagnelli, and G. Piscosquito. 2015. Mitochondrial dynamics and inherited peripheral nerve diseases. Neurosci Lett. 596:66–77. doi:10.1016/J.NEULET.2015.04.001.

Parsons, J.T., A.R. Horwitz, and M.A. Schwartz. 2010. Cell adhesion: integrating cytoskeletal dynamics and cellular tension. Nat Rev Mol Cell Biol. 11:633. doi:10.1038/NRM2957.

Parys, J.B., and H. de Smedt. 2012. Inositol 1,4,5-trisphosphate and its receptors. Adv Exp Med Biol. 740:255–279. doi:10.1007/978-94-007-2888-2_11.

Pelham, R.J., and Y.L. Wang. 1997. Cell locomotion and focal adhesions are regulated by substrate □ flexibility. Proc Natl Acad Sci U S A. 94:13661. doi:10.1073/PNAS.94.25.13661.

Penzes, P., M.E. Cahill, K.A. Jones, and D.P. Srivastava. 2008. Convergent CaMK and RacGEF signals control dendritic structure and function. Trends Cell Biol. 18:405–413. doi:10.1016/J.TCB.2008.07.002.

Pollard, T.D., and G.G. Borisy. 2003. Cellular motility driven by assembly and disassembly of actin filaments. Cell. 112:453–465. doi:10.1016/S0092-8674(03)00120-X.

Ridley, A.J., and A. Hall. 1992. The small GTP-binding protein rho regulates the assembly of focal adhesions and actin stress fibers in response to growth factors. Cell. 70:389–399. doi:10.1016/0092-8674(92)90163-7.

Rocha, A.G., A. Franco, A.M. Krezel, J.M. Rumsey, J.M. Alberti, W.C. Knight, N. Biris, E. Zacharioudakis, J.W. Janetka, R.H. Baloh, R.N. Kitsis, D. Mochly-Rosen, R.R. Townsend, E. Gavathiotis, and G.W. Dorn. 2018. MFN2 agonists reverse mitochondrial defects in preclinical models of Charcot-Marie-Tooth disease type 2A. Science. 360:336–341. doi:10.1126/SCIENCE.AAO1785.

Sabass, B., M.L. Gardel, C.M. Waterman, and U.S. Schwarz. 2008. High Resolution Traction Force Microscopy Based on Experimental and Computational Advances. Biophys J. 94:207. doi:10.1529/BIOPHYSJ.107.113670.

Sala, S., and P.W. Oakes. 2021. Stress fiber strain recognition by the LIM protein testin is cryptic and mediated by RhoA. Mol Biol Cell. 32:1758–1771. doi: 10.1091/MBC.E21-03-0156/MC-E21-03-0156-S10.MP4.

Saneyoshi, T., and Y. Hayashi. 2012. The Ca2+ and Rho GTPase signaling pathways underlying activity-dependent actin remodeling at dendritic spines. Cytoskeleton. 69:545–554. doi:10.1002/CM.21037.

Santel, A., and M.T. Fuller. 2001. Control of mitochondrial morphology by a human mitofusin. J Cell Sci. 114:867–874. doi:10.1242/jcs.114.5.867.

Seetharaman, S., and S. Etienne-Manneville. 2020. Cytoskeletal Crosstalk in Cell Migration. Trends Cell Biol. 30:720–735. doi:10.1016/J.TCB.2020.06.004.

Soderling, T.R. 1999. The Ca-calmodulin-dependent protein kinase cascade. Trends Biochem Sci. 24:232–236. doi:10.1016/S0968-0004(99)01383-3.

Strickland, A. v., A.P. Rebelo, F. Zhang, J. Price, B. Bolon, J.P. Silva, R. Wen, and S. Züchner. 2014. Characterization of the mitofusin 2 R94W mutation in a knock-in mouse model. J Peripher Nerv Syst. 19:152–164. doi:10.1111/JNS5.12066.

Stull, J.T., P.J. Lin, J.K. Krueger, J. Trewhella, and G. Zhi. 1998. Myosin light chain kinase: functional domains and structural motifs. Acta Physiol Scand. 164:471–482. doi:10.1111/J.1365-201X.1998.TB10699.X.

Sun, Y., R.N. Day, and A. Periasamy. 2011. Investigating protein-protein interactions in living cells using fluorescence lifetime imaging microscopy. Nature Protocols 2011 6:9.6:1324–1340. doi:10.1038/nprot.2011.364.

Takai, Y., T. Sasaki, K. Tanaka, and H. Nakanishi. 1995. Rho as a regulator of the cytoskeleton. Trends Biochem Sci. 20:227–231. doi:10.1016/S0968-0004(00)89022-2.

Tee, S.Y., J. Fu, C.S. Chen, and P.A. Janmey. 2011. Cell shape and substrate rigidity both regulate cell stiffness. Biophys J. 100. doi:10.1016/J.BPJ.2010.12.3744.

Tolias, K.F., J.B. Bikoff, A. Burette, S. Paradis, D. Harrar, S. Tavazoie, R.J. Weinberg, and M.E. Greenberg. 2005. The Rac1-GEF Tiam1 couples the NMDA receptor to the activity-dependent development of dendritic arbors and spines. Neuron. 45:525–538. doi:10.1016/J.NEURON.2005.01.024.

Totsukawa, G., Y. Yamakita, S. Yamashiro, D.J. Hartshorne, Y. Sasaki, and F. Matsumura. 2000. Distinct Roles of Rock (Rho-Kinase) and Mlck in Spatial Regulation of Mlc Phosphorylation for Assembly of Stress Fibers and Focal Adhesions in 3t3 Fibroblasts. J Cell Biol. 150:797. doi:10.1083/JCB.150.4.797.

Tsai, F.C., G.H. Kuo, S.W. Chang, and P.J. Tsai. 2015. Ca2+ signaling in cytoskeletal reorganization, cell migration, and cancer metastasis. Biomed Res Int. 2015. doi:10.1155/2015/409245.

Tsai, F.C., and T. Meyer. 2012. Ca2+ pulses control local cycles of lamellipodia retraction and adhesion along the front of migrating cells. Curr Biol. 22:837. doi:10.1016/J.CUB.2012.03.037.

Uehata, M., T. Ishizaki, H. Satoh, T. Ono, T. Kawahara, T. Morishita, H. Tamakawa, K. Yamagami, J. Inui, M. Maekawa, and S. Narumiya. 1997. Calcium sensitization of smooth muscle mediated by a Rho-associated protein kinase in hypertension. Nature 1997 389:6654. 389:990–994. doi:10.1038/40187.

Verhoeven, K., K.G. Claeys, S. Züchner, J.M. Schröder, J. Weis, C. Ceuterick, A. Jordanova, E. Nelis, E. de Vriendt, M. van Hul, P. Seeman, R. Mazanec, G.M. Saifi, K. Szigeti, P. Mancias, I. J. Butler, A. Kochanski, B. Ryniewicz, J. de Bleecker, P. van den Bergh, C. Verellen, R. van Coster, N. Goemans, M. Auer-Grumbach, W. Robberecht, V. Milic Rasic, Y. Nevo, I. Tournev, V. Guergueltcheva, F. Roelens, P. Vieregge, P. Vinci, M.T. Moreno, H.J. Christen, M.E. Shy, J.R. Lupski, J.M. Vance, P. de Jonghe, and V. Timmerman. 2006. MFN2 mutation distribution and genotype/phenotype correlation in Charcot-Marie-Tooth type 2. Brain. 129:2093–2102. doi:10.1093/BRAIN/AWL126.

Vicente-Manzanares, M., X. Ma, R.S. Adelstein, and A.R. Horwitz. 2009. Non-muscle myosin II takes centre stage in cell adhesion and migration. Nat Rev Mol Cell Biol. 10:778. doi:10.1038/NRM2786.

Vichai, V., and K. Kirtikara. 2006. Sulforhodamine B colorimetric assay for cytotoxicity screening. Nature Protocols 2006 1:3. 1:1112–1116. doi:10.1038/nprot.2006.179.

Wakatsuki, T., R.B. Wysolmerski, and E.L. Elson. 2003. Mechanics of cell spreading: role of myosin II. J Cell Sci. 116:1617–1625. doi:10.1242/JCS.00340.

Wang, X., B. Su, H.G. Lee, X. Li, G. Perry, M.A. Smith, and X. Zhu. 2009. Impaired balance of mitochondrial fission and fusion in Alzheimer’s disease. J Neurosci. 29:9090–9103. doi:10.1523/JNEUROSCI.1357-09.2009.

Wong, S.Y., T.A. Ulrich, L.P. Deleyrolle, J.L. MacKay, J.M.G. Lin, R.T. Martuscello, M.A. Jundi, B.A. Reynolds, and S. Kumar. 2015. Constitutive activation of myosin-dependent contractility sensitizes glioma tumor-initiating cells to mechanical inputs and reduces tissue invasion. Cancer Res. 75:1113–1122. doi:10.1158/0008-5472.CAN-13-3426.

Wynn, T.A. 2008. Cellular and molecular mechanisms of fibrosis. J Pathol. 214:199–210. doi:10.1002/PATH.2277.

Xie, Z., D.P. Srivastava, H. Photowala, L. Kai, M.E. Cahill, K.M. Woolfrey, C.Y. Shum, D.J. Surmeier, and P. Penzes. 2007. Kalirin-7 controls activity-dependent structural and functional plasticity of dendritic spines. Neuron. 56:640–656. doi:10.1016/J.NEURON.2007.10.005.

Xu, F., D. Ma, K.P. MacPherson, S. Liu, Y. Bu, Y. Wang, Y. Tang, C. Bi, T. Kwok, A.A. Chubykin, P. Yin, S. Calve, G.E. Landreth, and F. Huang. 2020. Three-dimensional nanoscopy of whole cells and tissues with in situ point spread function retrieval. Nature Methods 2020 17:5. 17:531–540. doi:10.1038/s41592-020-0816-x.

Ying, Z., F.R.C. Giachini, R.C. Tostes, and R.C. Webb. 2009. PYK2/PDZ-RhoGEF links Ca2+ signaling to RhoA. Arterioscler Thromb Vasc Biol. 29:1657. doi:10.1161/ATVBAHA.109.190892.

Zhao, J., J. Zhang, M. Yu, Y. Xie, Y. Huang, D.W. Wolff, P.W. Abel, and Y. Tu. 2012. Mitochondrial dynamics regulates migration and invasion of breast cancer cells. Oncogene 2013 32:40. 32:4814–4824. doi:10.1038/onc.2012.494.

Zhou, W., L. Cao, J. Jeffries, X. Zhu, C.J. Staiger, and Q. Deng. 2018. Neutrophil-specific knockout demonstrates a role for mitochondria in regulating neutrophil motility in zebrafish. DMM Disease Models and Mechanisms. 11. doi:10.1242/dmm.033027.

Zhou, W., A.Y. Hsu, Y. Wang, R. Syahirah, T. Wang, J. Jeffries, X. Wang, H. Mohammad, M.N. Seleem, D. Umulis, and Q. Deng. 2021. Mitofusin 2 regulates neutrophil adhesive migration and the actin cytoskeleton. J Cell Sci. 133. doi:10.1242/JCS.248880/VIDEO-11.

