## supplemental figures and movie legends for "Atypical peripheral actin band formation via overactivation of RhoA and Non-muscle myosin II in Mitofusin 2 deficient cells"

**Supplementary Figures**

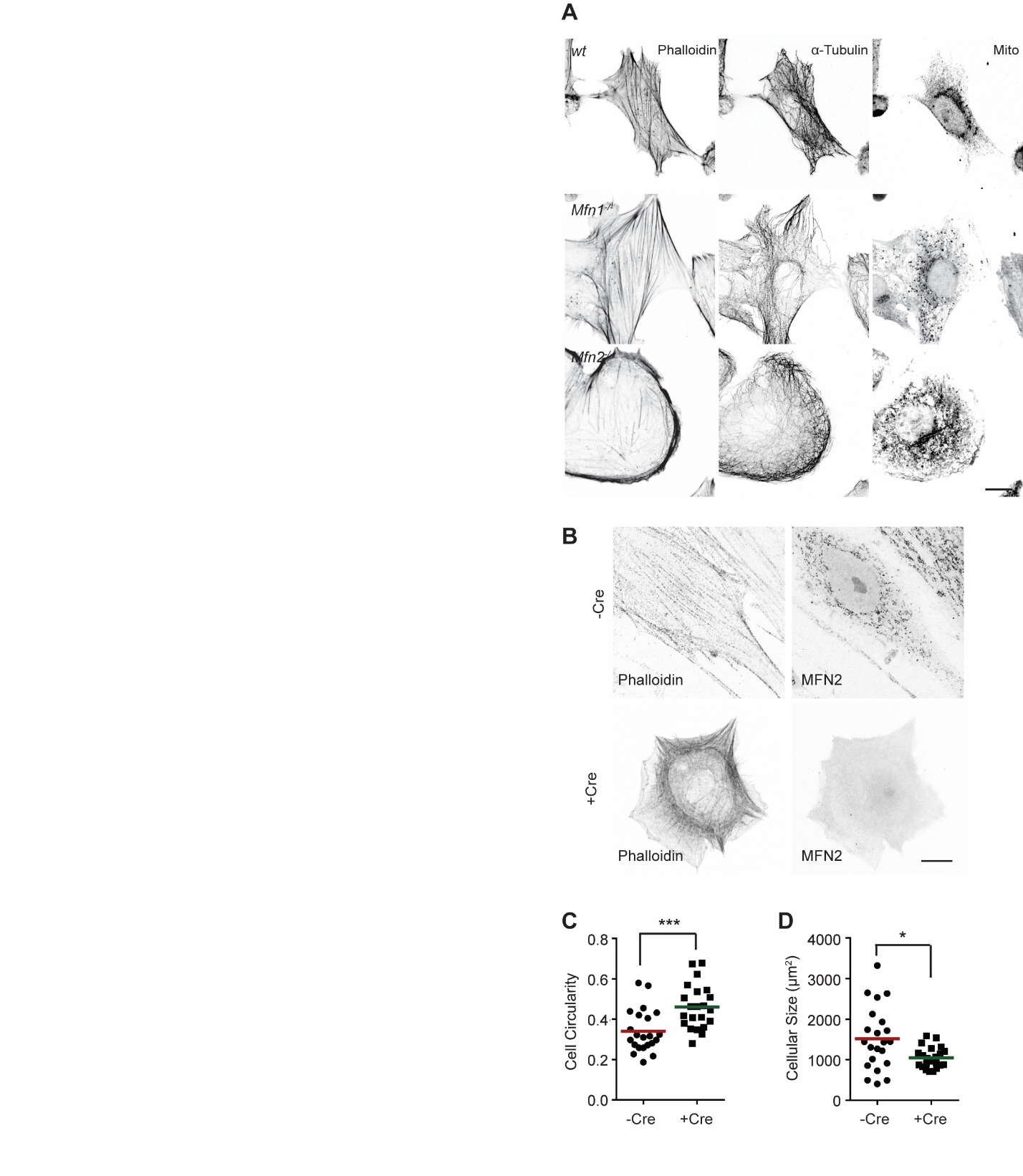

**Supplementary Figure 1. MFN2 deficiency changes MEF morphology.** (A) Immunofluorescence of F-actin (phalloidin), α-Tubulin, and Mitochondria (Mito Tracker) in wt, Mfn1-null, and Mfn2-null MEFs. (B-D) Cre-induced *Mfn2* disruption in MEFs from *Mfn2*^flox/flox^ mice display similar cell morphology as *Mfn2*-null MEFs (B). The individual points stand for the circularity (C) or size (D) of individual MEF cells. One representative result of three biological repeats is shown in A-B. Data are pooled from three independent experiments in C-D. n>25 cells are tracked and counted in C and D. ****P*≤0.001, *****P*<0.0001 (one-way ANOVA). Scale bars: 10µm.

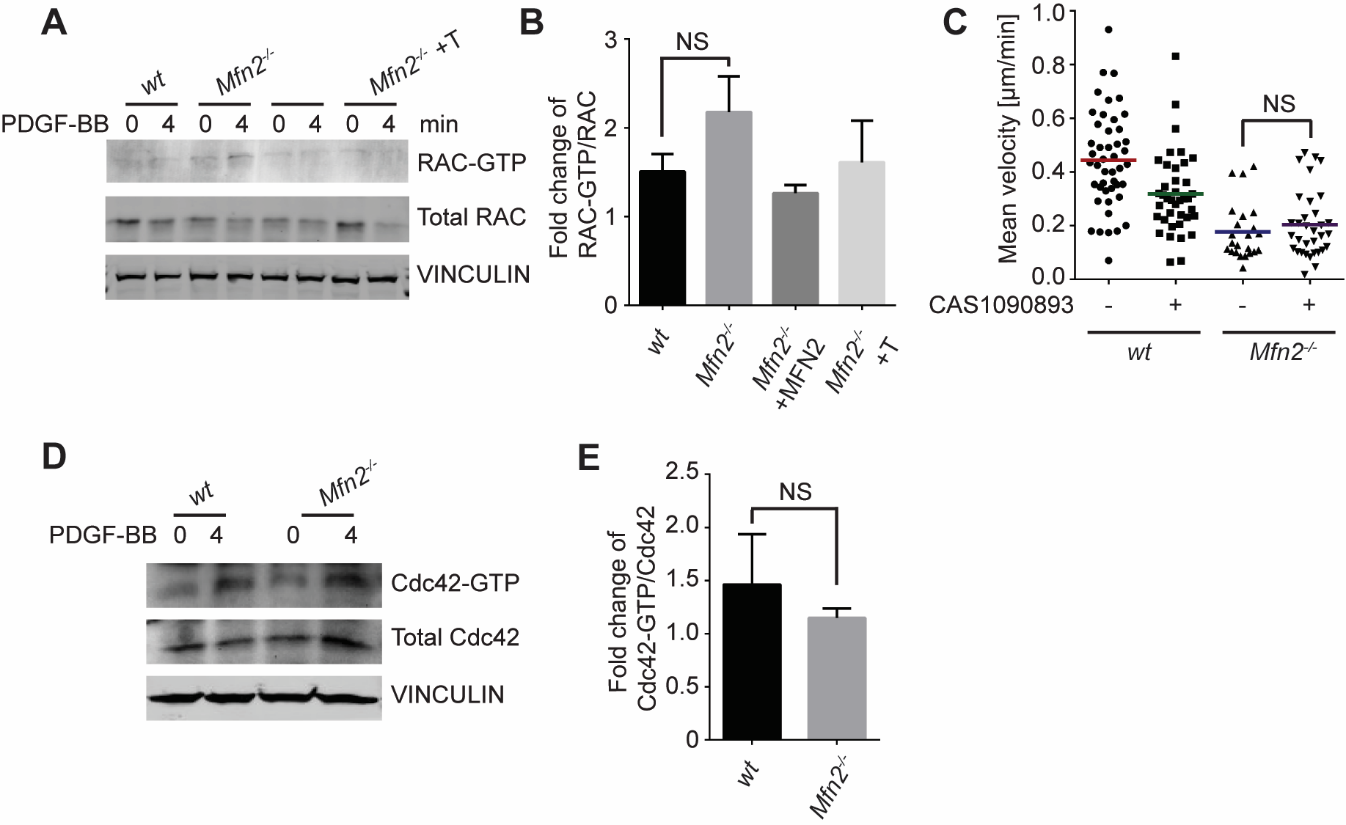
 **Supplementary Figure 2. Activities of Rac and Cdc42 are not increased in MFN2 deficient MEFs.** (A) Western blot and (B) quantification determining the amount of RAC-GTP total RAC protein in *wt*, *Mfn2*-null MEFs, *Mfn2*-null MEFs with MFN2 re-expressed, or with an artificial ER-mitochondria tether. The indicated cell lines were treated with 25ng/ml PDGF-BB for the indicated time. (C) Quantification of the velocity of indicated MEF cells treated with DMSO or Rac inhibitor CAS1090893. (D, E) Cdc42 pulldown activation assay demonstrates no difference in Cdc42-GTP in *wt and Mfn2*-null MEFs. (D) Western blot and (E) quantification determining the amount of Cdc42-GTP total Cdc42 protein in *wt*, *Mfn2*-null MEFs treated with 25ng/ml PDGF-BB for the indicated time. One representative result of three biological repeats in A, D. Data are pooled from three independent experiments in B, C, E. Error bars represent s.d. NS, non-significant [two-way ANOVA (C), one-way ANOVA (B), unpaired t-test (E)].

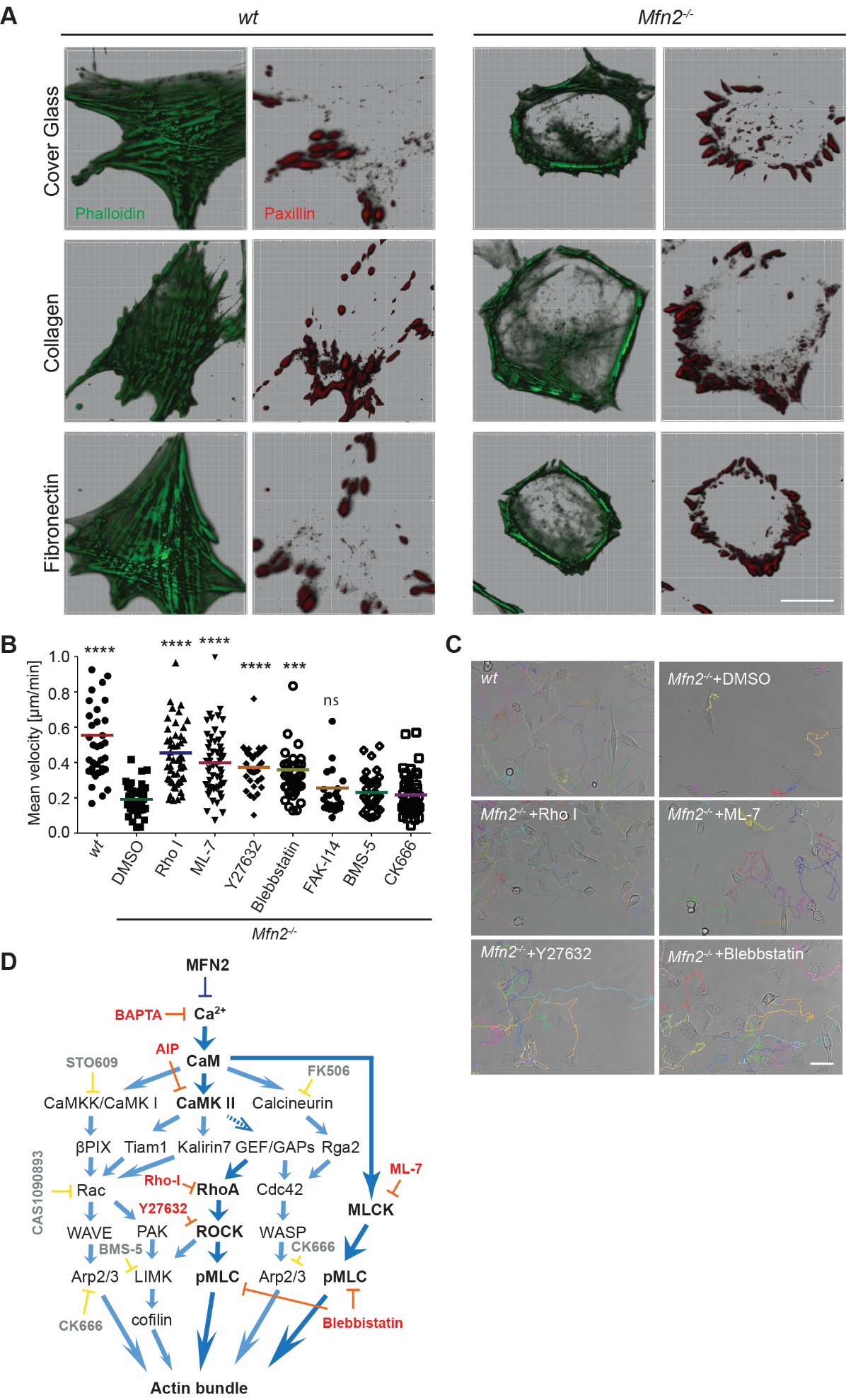

**Supplementary Figure 3. “PAB” structure in MFN2 deficient MEFs is not affected by the surface coating.** (A) Immunofluorescence of F-actin (phalloidin) and Paxillin in *wt* and Mfn2-null MEFs cultured on uncoated, collagen-coated, or fibrinogen-coated cover glasses. (B) Velocity quantification of indicated MEF cells treated with DMSO, the Rho inhibitor-I, the MLCK inhibitor ML-7, the ROCK inhibitor Y27632, the myosin inhibitor Blebbstatin, FAK inhibitor-14, the LIM kinase inhibitor BMS-5, or the Arp2/3 inhibitor CK666.(C) Representative images with individual tracks of indicated MEF cells treated with DMSO or indicated inhibitors. (D) A small molecular inhibitor screen revealed the schematic of the effectors and their inhibitors (red) in MFN2 regulated signaling network leading to Actin bundle. Data are pooled from three independent experiments in B and *n*>30 cells are quantified. ****P*≤0.001, *****P*<0.0001 (one-way ANOVA comparing each group to the average of *Mfn2^-/-^* DMSO group). Scale bars: 50 µm.

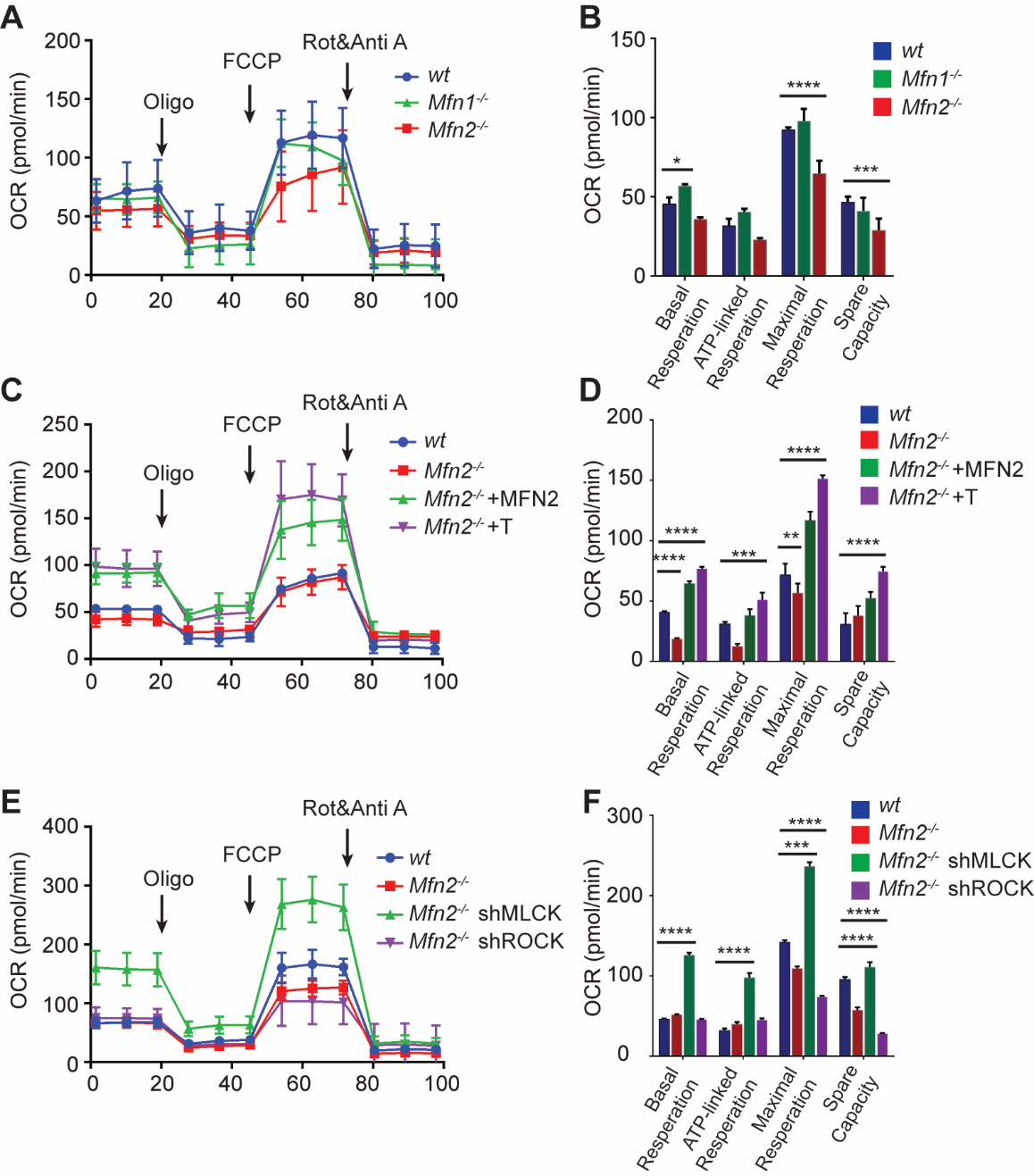

**Supplementary Figure 4. The rescue cell lines with restored motility show different oxygen consumption rates.** (A, C, E) Representative graph showing oxygen consumption rate (OCR) of indicated MEF cells. (B, D, F) Basal respiration, ATP-linked respiration, maximal respiratory, and spare capacity of indicated cell lines. One representative result of three biological repeats in A-F E. Error bars represent s.d. **P*≤0.05, ***P*≤0.01, ****P*≤0.001, *****P*<0.0001 (two-way ANOVA).

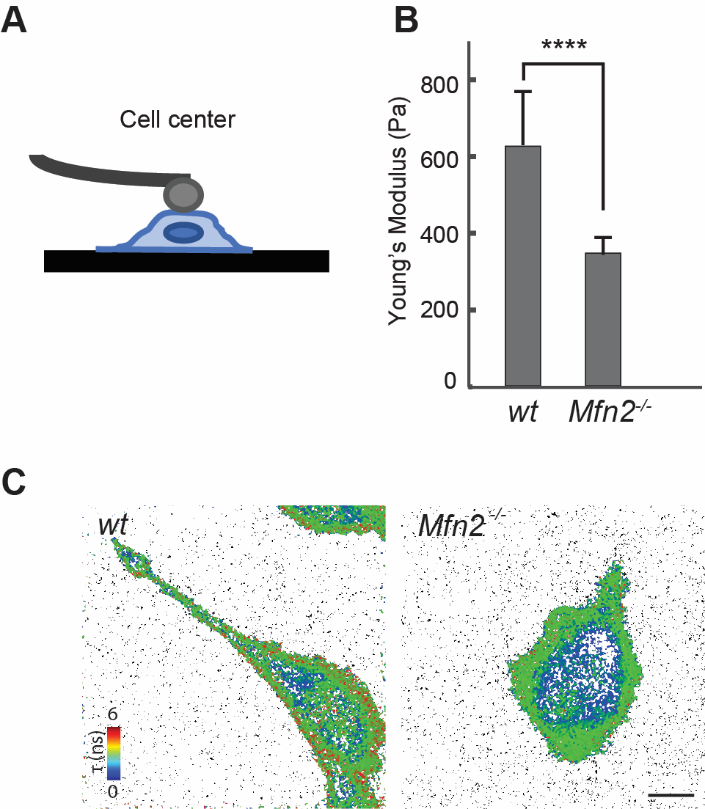

**Supplementary Figure 5. *Mfn2*-null MEFs exhibit altered cell stiffness and membrane tension.** (A) Schematic of the cantilever probe indenting cells at cell center during AFM process. (B) Measurement of effective cell modulus (Mean ± SEM) of *wt* or *Mfn2*-null MEFs (n ≥ 15 for each group). (C) Representative FLIM images showing the lifetime of the Flipper-TR probe on *wt* and *Mfn2*-null MEFs. One representative result of three biological repeats in A-C. Data are pooled from three independent experiments in B. Error bars represent s.d. *****P*<0.0001 [unpaired t-test (B)].

**Supplementary Movie legends**

**Movie 1. MFN2 regulates the migration and spreading of MEFs.**

Cell spreading and random migration of wt, Mfn2-null, and Mfn1-null MEFs in the µ-slide 15 mins after plating. Time-lapse images were taken every ten minutes for 16 hours and 40 minutes. Individual MEFs were tracked for velocity quantification. Scale bar: 50m.

**Movie 2. MFN2 re-expression, but not MFN1, restore migratory defects in Mfn2-null MEFs.**

Cell spreading and random migration of wt and Mfn2-null MEFs with *vec*, MFN1, or MFN2 re-expressed in the µ-slide. Time-lapse images were taken every ten minutes for 17 hours and 50 minutes. Individual MEFs were tracked for velocity quantification. Scale bar: 50m.

**Movie 3. Cytosolic Ca^2+^ inhibition restores the migration defects in Mfn2-null MEF cells.**

Cell spreading and random migration of wt and Mfn2-null MEFs treated with DMSO or BAPTA-AM (20 μM) in the μ-slide. Time-lapse images were taken every ten minutes for 14 hours and 30 minutes. MEFs were tracked for velocity quantification. Scale bar: 50 μm.

**Movie 4. Restoring the ER-mitochondrial tether rescues the migration defects in Mfn2-null MEF cells.**

Cell spreading and random migration of wt, Mfn2-null MEFs with *vec* or with synthetic tether construct in the µ-slide. Time-lapse images were taken every ten minutes for 17 hours and 50 minutes. Individual MEFs were tracked for velocity quantification. Scale bar: 50m.

**Movie 5. CaMKII inhibition rescues the migration defects in Mfn2-null MEF cells.**

Cell spreading and random migration of wt and Mfn2-null MEFs treated with DMSO or AIP (40 μM) in the μ-slide. Time-lapse images were taken every ten minutes for 17 hours and 50 minutes. MEFs were tracked for velocity quantification. Scale bar: 50 μm.

**Movie 6. Expression of CaMKII-DN rescues the migration defects in Mfn2-null MEF cells.**

Cell spreading and random migration of wt, Mfn2-null MEFs with *vec*, CaMKII-WT, or CaMKII-DN in the μ-slide. Time-lapse images were taken every ten minutes for 17 hours and 50 minutes. MEFs were tracked for velocity quantification. Scale bar: 50 μm.

**Movie 7. Small molecule inhibitors targeting RhoA and MLCK downstream signaling pathways rescue the migration defects in Mfn2-null MEF cells.**

Cell spreading and random migration of wt treated with DMSO and Mfn2-null MEFs treated with DMSO, RhoA inhibitor-I (0.1µg/ml), ML-7 (2µM), Y29632 (5µM) or Blebbistatin (4µM) in the μ-slide. Time-lapse images were taken every ten minutes for 14 hours and 30 minutes. MEFs were tracked for velocity quantification. Scale bar: 50 μm.
